# Itaconate is metabolized to 2-hydroxymethylsuccinate through a CoA-independent degradation pathway in mitochondria

**DOI:** 10.1101/2025.09.29.679238

**Authors:** Fangfang Chen, Laura Hadam, Birte Dowerg, Vadim S. Korotkov, Connor S.R. Jankowski, Ulrike Beutling, Hanna F. Willenbockel, Ulrike Brandt, Eva Medina, Andre Fleissner, Henrikus Garritsen, Michael Jarek, Raimo Franke, Thomas Weichhart, Mark Brönstrup, Meina Neumann-Schaal, Thekla Cordes

**Affiliations:** Department of Bioinformatics and Biochemistry, Technische Universität Braunschweig, Rebenring 56, 38106 Braunschweig, Germany; Cellular Metabolism in Infection Research Group, Helmholtz Centre for Infection Research, Inhoffenstraße 7, 38124 Braunschweig, Germany; Department of Chemical Biology, Helmholtz Centre for Infection Research, Inhoffenstraße 7, 38124 Braunschweig, Germany; Institute of Medical Genetics, Center for Pathobiochemistry and Genetics, Medical University of Vienna, Vienna, Austria; Institute of Genetics, Technische Universität Braunschweig, Braunschweig, Germany; Infection Immunology Research Group, Helmholtz Centre for Infection Research, Inhoffenstraße 7, 38124 Braunschweig, Germany; Institute of Transfusion Medicine, Klinikum Braunschweig, Braunschweig, Germany; Genome Analytics, Helmholtz Centre for Infection Research, Inhoffenstraße 7, 38124 Braunschweig, Germany; Leibniz Institute DSMZ–German Collection of Microorganisms and Cell Cultures, Department of Metabolomics and Services, 38124 Braunschweig, Germany; Braunschweig Integrated Centre of Systems Biology (BRICS), Braunschweig, Germany

**Keywords:** itaconate, mitochondrial metabolism, tracing, mass spectrometry, degradation, C_5_ dicarboxylate pathway, 2-hydroxymethylsuccinate (2HMS), itamalate

## Abstract

The immunometabolite itaconate modulates cellular metabolism and is converted into structurally similar C_5_ dicarboxylates that require advanced analytics to decipher their metabolic fate. Here, we employ high-resolution mass spectrometry and tracing approaches and identify 2-hydroxymethylsuccinate (2HMS) as a previously unrecognized C_5_ dicarboxylate derived from itaconate. 2HMS synthesis occurs during inflammatory responses and upon itaconate treatment, as detected by ^13^C itaconate tracing. Pathway analysis reveals that methylglutaconyl-CoA hydratase (AUH) drives 2HMS synthesis through a CoA-independent conversion (CIC) pathway. This pathway is distinct from the CoA-dependent conversion (CDC) pathway that generates mesaconate and itaconyl-CoA influencing B_12_-dependent processes. *In vivo* inflammation studies reveal that adipose tissue prefers CIC to produce 2HMS and liver favors CDC-mediated mesaconate synthesis, highlighting tissue-specific itaconate degradation routes. This study identifies a new branch of itaconate metabolism, provides an analytical framework to resolve C_5_ dicarboxylate networks, and links 2HMS to inflammation and mitochondrial metabolism that might be targeted therapeutically.

## Introduction

Metabolites have emerged as signaling molecules and their dynamics influence cellular processes. Among these, oncometabolites and immunometabolites are modulators of the cellular homeostasis influencing disease progression and outcome^1^. For instance, the five-carbon (C_5_) dicarboxylate 2-hydroxyglutarate (2HG) is implicated in cancer and inflammation^2–4^, while pro-inflammatory conditions drive the synthesis of the immunomodulatory metabolite itaconate^5,6^. Metabolites that exert biological effects demand precise analysis of their enzymatic synthesis and degradation pathways to establish their roles in disease mechanisms.

Itaconate accumulates to millimolar levels in macrophages and is synthesized via immune-responsive gene 1 protein (IRG1), also known as *cis*-aconitate decarboxylase (ACOD1), encoded by immunoresponsive gene 1 (*Irg1*)^7,8^. IRG1 levels are highly increased in macrophages in response to inflammatory stimuli linking metabolism to immune defenses. For instance, mitochondrial transporters and nitric oxide signaling modulate itaconate levels, thereby fine-tuning the effects of itaconate on cellular metabolism and macrophage function^9,10^. Further, itaconate transiently inhibits succinate dehydrogenase (SDH) activity^11,12^ and influences methylmalonyl-CoA mutase (MUT) activity^13,14,30^ impacting tricarboxylic acid (TCA) cycle flux, Coenzyme A (CoA) homeostasis^14,15^, and amino acid and lipid metabolism^16–19^. This metabolic rewiring enables itaconate to influence bacterial infections^7,20^, immune signaling^21–24^, and tumor growth^1,25–27^. Itaconate is rapidly cleared from plasma resulting in a dynamic regulation of metabolites such as succinate^16,28^. However, its degradation pathways remain incompletely understood and, thus, deciphering the fate of itaconate may advance our understanding of biological processes associated with itaconate.

Our recent ^13^C itaconate tracing study revealed that itaconate is primarily cleared via the kidneys and that a minor fraction fuels TCA cycle metabolism^28^. Itaconate is converted to itaconyl-CoA catalyzed by succinyl-CoA synthetase (SCS), thereby influencing the subsequent substrate-level phosphorylation within the TCA cycle^15,29,30^. This dynamic turnover highlights the multifaceted effects of itaconate whose function derives not only from the parent compound but also from its degradation products, such as citramalate and mesaconate ^28,31–33^. However, the full spectrum of itaconate-derived metabolites and their tissue-specific metabolism remains poorly understood. Further, the structural similarity among C_5_ dicarboxylates including itaconate, mesaconate, citramalate, 3-methylmalate, 3-hydroxyglutarate (3HG), and 2HG complicates their discrimination by mass spectrometry and requires advanced analytical approaches to prevent misidentification^34^.

Here, we applied high-resolution mass spectrometry and tracing experiments to characterize the fate of itaconate and to identify potential conversion pathways. Our analytical approach reliably distinguishes C_5_ dicarboxylates including itaconate, mesaconate, 2HG, 3HG, and citramalate. Of note, we identified 2-hydroxymethylsuccinate (2HMS, also known as itamalate) as a previously uncharacterized C_5_ dicarboxylate metabolite derived from itaconate which is synthesized through a CoA-independent pathway catalyzed by mitochondrial methylglutaconyl-CoA hydratase (AUH, also known as AU RNA binding protein/enoyl-CoA hydratase). Our findings define a new branch of itaconate metabolism, provide tools to dissect C_5_ dicarboxylate complexity, and identify 2HMS and itaconate pathways as potential targets with therapeutic implications.

## Results

### Discovery of 2-hydroxymetylsuccinate (2HMS) in stimulated macrophages

Itaconate is synthesized *de novo* from the TCA cycle intermediate cis-aconitate via ACOD1 in macrophages under pro-inflammatory conditions^7^. To investigate the fate of itaconate and metabolic reprogramming with induced ACOD1 activity, we quantified metabolism over time in LPS-stimulated RAW264.7 macrophage-like cells. We observed that cells produced high amounts of itaconate and mesaconate peaking at 6 and 12 h post-stimulation, respectively, and then declining indicating potential degradation (Fig. 1a). Of note, our mass spectrometry data identified the presence of a previously uncharacterized metabolite that accumulated in response to LPS treatment or in cultures with exogenous itaconate (Fig. 1a). While itaconate and mesaconate levels declined by 24 h, levels of this uncharacterized metabolite remained elevated suggesting a distinct metabolic fate compared to mesaconate synthesis.

**Figure 1:**
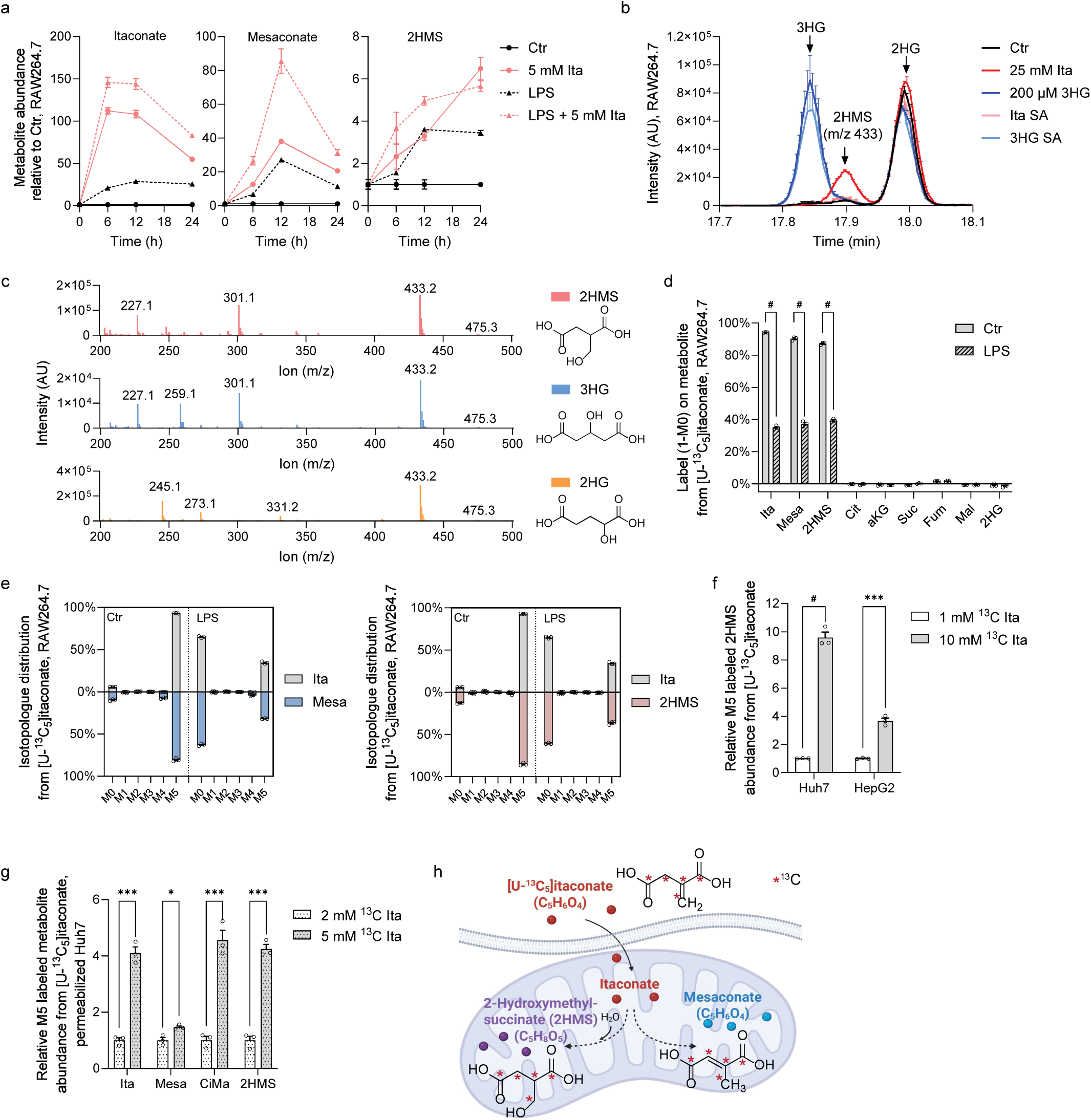
^13^C itaconate is converted into the C_5_ dicarboxylate metabolite 2-hydroxymethylsuccinate (2HMS). **a,** Metabolite abundance of itaconate, mesaconate, and 2-hydroxymethylsuccinate (2HMS) in RAW264.7 cells cultured with 10 ng/ml LPS and 5 mM itaconate for 6, 12, and 24 h. **b,** GC-MS Ion (m/z 433) chromatogram depicting 2HMS, 2-hydroxyglutarate (2HG), and 3-hydroxyglutarate (3HG) in RAW264.7 cells with standard addition (SA) or cultured with 25 mM itaconate and 200 µM 3HG for 24 h. **c,** GC-MS Total Ions Chromatograms (TIC) and chemical structures of 3HG, 2HG, and 2HMS. **d,** Label on metabolites in RAW264.7 cells cultured with 1 mM [U-^13^C_5_]itaconate and 10 ng/ml LPS for 24 h compared to control condition. **e,** Isotopologue distribution of mesaconate (blue) and 2HMS (red) compared to itaconate (grey) in RAW264.7 cells cultured with 1 mM [U-^13^C_5_]itaconate and 10 ng/ml LPS for 24 h compared to control condition. **f,** Labeled 2HMS abundance in Huh7 and HepG2 cells cultured for 24 h with 1 and 10 mM [U-^13^C_5_]itaconate. **g,** Labeled mitochondrial metabolite abundances in permeabilized Huh7 cells cultured with 2 and 5 mM [U-^13^C_5_]itaconate. **h,** Schematic depicting the synthesis of 2HMS and mesaconate from ^13^C itaconate in mitochondria prepared with Biorender. Data are presented as means ± s.e.m. obtained from n = 3 cellular replicates. Similar results were obtained from three (b, c, d, e) or two (a, f, g) independent experiments. *P* values were calculated by multiple unpaired *t*-test with * *p* < 0.05, ** *p* < 0.01, *** *p* < 0.001, ^#^ *p* < 0.0001.

To gain insights into the chemical structure of the newly detected metabolite, we analyzed high-resolution gas chromatography (GC) coupled to tandem mass spectrometry (MS/MS) data obtained for silylated compounds and observed that the metabolite had a retention time of 18.16 min and a characteristic fragment ion m/z 433 (Table S1). We assigned structural formulas to characteristic fragments indicating that 2HMS contains five carbons with the molecular formula C_5_H_8_O_5_. The latter is common for diverse C_5_ dicarboxylate compounds including citramalate (CiMa), 2-hydroxyglutarate (2HG), 3-hydroxyglutarate (3HG), 3-methylmalate (3MeMa), or 2-hydroxymethylsuccinate (2HMS) (Table S1). To reliably distinguish these compounds based on retention times, we analyzed synthetic standards. Since a synthetic 2HMS standard was unavailable, we established a chemical synthesis by alkaline hydrolysis of commercially available 5-oxotetrahydrofuran-3-carboxylic acid (Fig. S1a). The molecular structure of the 2HMS compound was unequivocally confirmed by 1D- and 2D-nuclear magnetic resonance (NMR) spectroscopy (Fig. S1b-f). Importantly, standard addition (SA) experiments led to a co-elution of the synthetic 2HMS standard with the endogenously produced uncharacterized metabolite in cultures with itaconate (Fig. S1g). A comparison of retention times of these C_5_ dicarboxylates clearly differentiated CiMa and 3MeMa from the newly detected metabolite (Fig. S2a). The mass spectra of these C_5_ dicarboxylates depict that 3HG, 2HG, citramalate, 3MeMa, and 2HMS share the same m/z 433 fragment ion, but all have a unique mass spectrum with characteristic ions to distinguish these compounds (Table S1, Fig. S2b). Further, MS2 high-resolution mass spectra from the parent ion m/z = 433 depict further differences between 3HG, 2HG, and the newly detected metabolite (Fig. S2c).

Thus, our analytical GC-MS approach distinguishes 2HMS from other C_5_ dicarboxylate compounds, specifically 2HG and 3HG which all share the characteristic m/z 433 fragment ion, based on their characteristic fragmentation patterns and retention times (Fig. 1b, c, Table S1). Itaconate treatment markedly increased 2HMS abundance, whereas 2HG and 3HG levels were less affected (Fig. 1b). 2HMS was detected exclusively in itaconate-treated cultures, and our control experiments excluded spontaneous chemical degradation of itaconate or 3HG as its source (Fig. 1b, Fig. S1g). We therefore discovered that LPS-activated macrophages, or cultured with exogenous itaconate, synthesize the previously uncharacterized metabolite 2-hydroxymethylsuccinate (2HMS), also known as itamalate.

Finally, we purified a metabolite fraction containing endogenously synthesized 2HMS from RAW264.7 cells cultured in the presence of itaconate and performed high-resolution liquid chromatography-tandem mass spectroscopy (LC-MS/MS) analysis. Exact mass measurements proposed the sum formula C_5_H_8_O_5,_ which is characteristic of several structurally similar compounds (Table 1). Our comparative profiling further demonstrates overlapping elution times among C_5_ dicarboxylates that complicated computational identification in LC-MS analysis (Fig. S3a). To resolve this, we determined high resolution MS1 and MS2 mass spectra for the C_5_ dicarboxylates and identified distinct precursor ions (m/z 129.0195 for itaconate; m/z 147.0296 for 2HMS, 2HG, 3HG, citramalate, and 3-methylmalate) and fragmentation patterns for each C_5_ dicarboxylate compound (Fig. S3b). The isolated compound fraction from itaconate-treated cells at RT 1.82 min contained 2HG, while the fraction at RT 1.7 min contained 2HMS confirming the presence of 2HMS in culture conditions. Of note, the 2HMS anion exhibited two characteristic fragment ions at m/z = 117 and m/z 99 due to the losses of [C,H_2_,O] and [C,H_4_,O_2_], respectively (Fig. S3b). They may be derived from a retro-aldol cleavage in the gas phase and are therefore not found in any of the other isomers. Collectively, our mass spectrometry approaches combined with synthetic standard validation identified 2HMS as a previously overlooked, endogenously synthesized compound in the C_5_ dicarboxylate metabolic network.

**Table 1:**
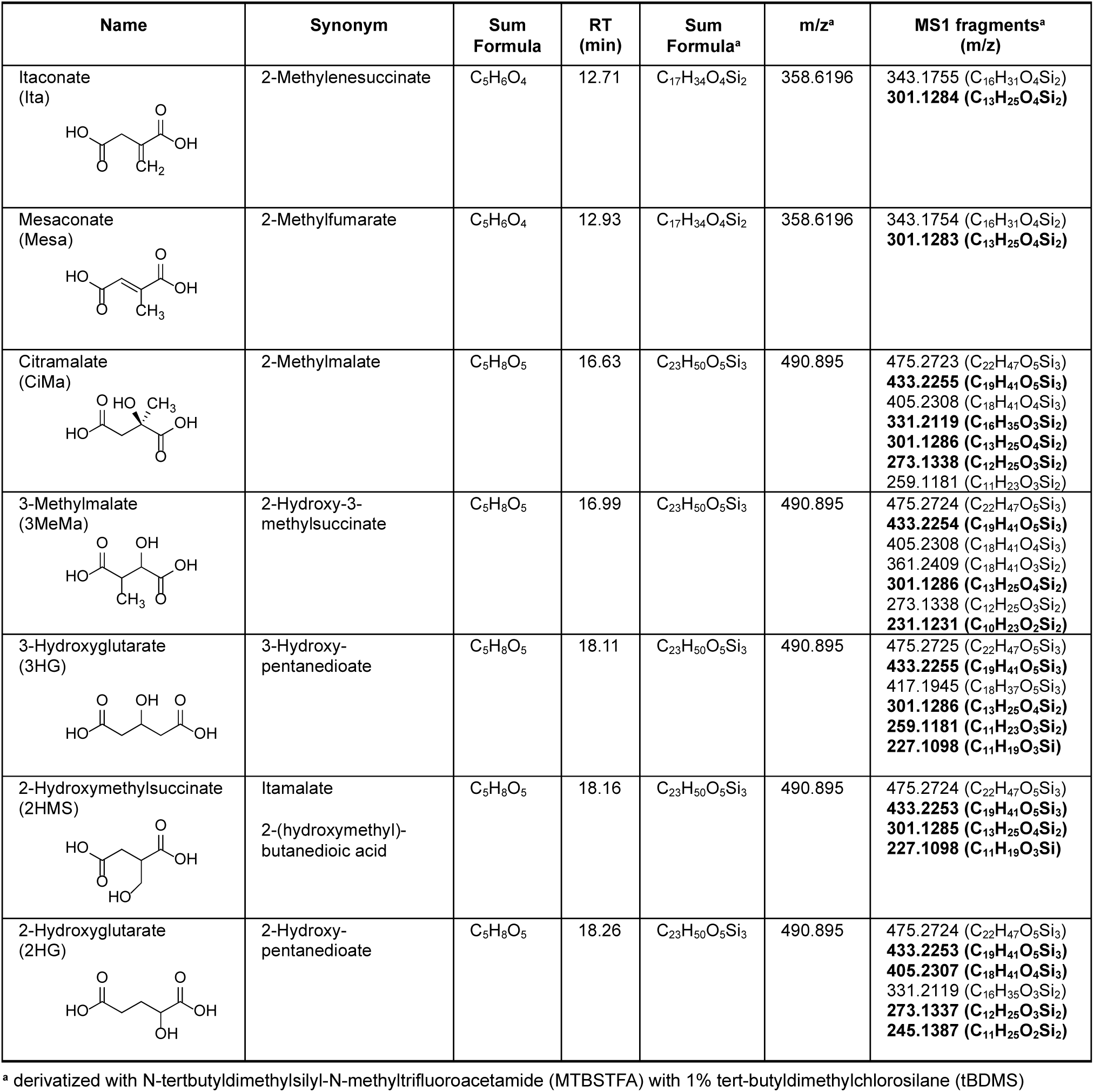
Chemical properties and mass fragmentation to distinguish C_5_ dicarboxylate compounds via GC-MS/MS. Chemical information and characteristic ions of derivatized fragments (MS1) using N-tertbutyldimethylsilyl-N-methyltrifluoroacetamide (MTBSTFA) with 1% tert-butyldimethylchlorosilane (tBDMS). MS1 fragments highlighted in bold are used for compound identification.

| Name | Synonym | Sum Formula | RT (min) | Sum Formula<sup>a</sup> | m/z<sup>a</sup> | MS1 fragments<sup>a</sup> (m/z) |
| --- | --- | --- | --- | --- | --- | --- |
| Itaconate<br>(Ita) | 2-Methylenesuccinate | C<sub>5</sub>H<sub>6</sub>O<sub>4</sub> | 12.71 | C<sub>17</sub>H<sub>34</sub>O<sub>4</sub>Si<sub>2</sub> | 358.6196 | 343.1755 (C<sub>16</sub>H<sub>31</sub>O<sub>4</sub>Si<sub>2</sub>)<br>**301.1284 (C<sub>13</sub>H<sub>25</sub>O<sub>4</sub>Si<sub>2</sub>)** |
| Mesaconate<br>(Mesa) | 2-Methylfumarate | C<sub>5</sub>H<sub>6</sub>O<sub>4</sub> | 12.93 | C<sub>17</sub>H<sub>34</sub>O<sub>4</sub>Si<sub>2</sub> | 358.6196 | 343.1754 (C<sub>16</sub>H<sub>31</sub>O<sub>4</sub>Si<sub>2</sub>)<br>**301.1283 (C<sub>13</sub>H<sub>25</sub>O<sub>4</sub>Si<sub>2</sub>)** |
| Citramalate<br>(CiMa) | 2-Methylmalate | C<sub>5</sub>H<sub>8</sub>O<sub>5</sub> | 16.63 | C<sub>23</sub>H<sub>50</sub>O<sub>5</sub>Si<sub>3</sub> | 490.895 | 475.2723 (C<sub>22</sub>H<sub>47</sub>O<sub>5</sub>Si<sub>3</sub>)<br>**433.2255 (C<sub>19</sub>H<sub>41</sub>O<sub>5</sub>Si<sub>3</sub>)**<br>405.2308 (C<sub>18</sub>H<sub>41</sub>O<sub>4</sub>Si<sub>3</sub>)<br>**331.2119 (C<sub>16</sub>H<sub>35</sub>O<sub>3</sub>Si<sub>2</sub>)**<br>**301.1286 (C<sub>13</sub>H<sub>25</sub>O<sub>4</sub>Si<sub>2</sub>)**<br>**273.1338 (C<sub>12</sub>H<sub>25</sub>O<sub>3</sub>Si<sub>2</sub>)**<br>259.1181 (C<sub>11</sub>H<sub>23</sub>O<sub>3</sub>Si<sub>2</sub>) |
| 3-Methylmalate<br>(3MeMa) | 2-Hydroxy-3-methylsuccinate | C<sub>5</sub>H<sub>8</sub>O<sub>5</sub> | 16.99 | C<sub>23</sub>H<sub>50</sub>O<sub>5</sub>Si<sub>3</sub> | 490.895 | 475.2724 (C<sub>22</sub>H<sub>47</sub>O<sub>5</sub>Si<sub>3</sub>)<br>**433.2254 (C<sub>19</sub>H<sub>41</sub>O<sub>5</sub>Si<sub>3</sub>)**<br>405.2308 (C<sub>18</sub>H<sub>41</sub>O<sub>4</sub>Si<sub>3</sub>)<br>361.2409 (C<sub>18</sub>H<sub>41</sub>O<sub>3</sub>Si<sub>2</sub>)<br>**301.1286 (C<sub>13</sub>H<sub>25</sub>O<sub>4</sub>Si<sub>2</sub>)**<br>273.1338 (C<sub>12</sub>H<sub>25</sub>O<sub>3</sub>Si<sub>2</sub>)<br>**231.1231 (C<sub>10</sub>H<sub>23</sub>O<sub>2</sub>Si<sub>2</sub>)** |
| 3-Hydroxyglutarate<br>(3HG) | 3-Hydroxy-pentanedioate | C<sub>5</sub>H<sub>8</sub>O<sub>5</sub> | 18.11 | C<sub>23</sub>H<sub>50</sub>O<sub>5</sub>Si<sub>3</sub> | 490.895 | 475.2725 (C<sub>22</sub>H<sub>47</sub>O<sub>5</sub>Si<sub>3</sub>)<br>**433.2255 (C<sub>19</sub>H<sub>41</sub>O<sub>5</sub>Si<sub>3</sub>)**<br>417.1945 (C<sub>18</sub>H<sub>37</sub>O<sub>5</sub>Si<sub>3</sub>)<br>**301.1286 (C<sub>13</sub>H<sub>25</sub>O<sub>4</sub>Si<sub>2</sub>)**<br>**259.1181 (C<sub>11</sub>H<sub>23</sub>O<sub>3</sub>Si<sub>2</sub>)**<br>**227.1098 (C<sub>11</sub>H<sub>19</sub>O<sub>3</sub>Si)** |
| 2-Hydroxymethylsuccinate<br>(2HMS) | Itamalate<br>2-(hydroxymethyl)-butanedioic acid | C<sub>5</sub>H<sub>8</sub>O<sub>5</sub> | 18.16 | C<sub>23</sub>H<sub>50</sub>O<sub>5</sub>Si<sub>3</sub> | 490.895 | 475.2724 (C<sub>22</sub>H<sub>47</sub>O<sub>5</sub>Si<sub>3</sub>)<br>**433.2253 (C<sub>19</sub>H<sub>41</sub>O<sub>5</sub>Si<sub>3</sub>)**<br>**301.1285 (C<sub>13</sub>H<sub>25</sub>O<sub>4</sub>Si<sub>2</sub>)**<br>**227.1098 (C<sub>11</sub>H<sub>19</sub>O<sub>3</sub>Si)** |
| 2-Hydroxyglutarate<br>(2HG) | 2-Hydroxy-pentanedioate | C<sub>5</sub>H<sub>8</sub>O<sub>5</sub> | 18.26 | C<sub>23</sub>H<sub>50</sub>O<sub>5</sub>Si<sub>3</sub> | 490.895 | 475.2724 (C<sub>22</sub>H<sub>47</sub>O<sub>5</sub>Si<sub>3</sub>)<br>**433.2253 (C<sub>19</sub>H<sub>41</sub>O<sub>5</sub>Si<sub>3</sub>)**<br>**405.2307 (C<sub>18</sub>H<sub>41</sub>O<sub>4</sub>Si<sub>3</sub>)**<br>331.2119 (C<sub>16</sub>H<sub>35</sub>O<sub>3</sub>Si<sub>2</sub>)<br>**273.1337 (C<sub>12</sub>H<sub>25</sub>O<sub>3</sub>Si<sub>2</sub>)**<br>**245.1387 (C<sub>11</sub>H<sub>25</sub>O<sub>2</sub>Si<sub>2</sub>)** |

<sup>a</sup> derivatized with N-tertbutyldimethylsilyl-N-methyltrifluoroacetamide (MTBSTFA) with 1% tert-butyldimethylchlorosilane (tBDMS)

### Itaconate is converted into the C_5_ dicarboxylate metabolite 2-hydroxymethylsuccinate (2HMS)

2HMS is a five-carbon metabolite whose synthesis depends on itaconate, indicating a potential precursor–product relationship (Fig. 1a). To identify whether 2HMS is directly derived from itaconate, we cultured RAW264.7 macrophage-like cells in the presence of uniformly (U) labeled [U-^13^C_5_]itaconate. Itaconate was highly labeled in control conditions indicating minimal endogenous production. LPS-stimulation decreased labeling on itaconate due to *de novo* synthesis of unlabeled itaconate from cellular carbon sources (Fig. 1d). We also detected labeled mesaconate with a similar labeling pattern compared to itaconate while other TCA cycle intermediates and 2HG were not labeled. This data confirmed that itaconate is metabolized to mesaconate and that itaconate does not label TCA cycle intermediates in our culture system^15,16,28,32^.

Further, we observed that the newly identified metabolite 2HMS was highly labeled from ^13^C itaconate. Of note, labeling decreased upon LPS treatment indicating *de novo* synthesis from itaconate similar as observed for mesaconate (Fig. 1d). The mass spectrum of 2HMS depicted a characteristic fragment ion at m/z 433 with strong M5 (m/z 438) labeling from [U-^13^C_5_]itaconate that closely aligned with that of itaconate and mesaconate (Fig. 1e). This tracing data confirms that 2HMS is derived from itaconate *de novo* and retains the full C_5_ carbon backbone of labeled itaconate.

Previous studies identified the liver as a key site for itaconate degradation^28,31^. Therefore, we traced hepatocyte cultures with [U-^13^C_5_]itaconate and detected M5 labeled 2HMS confirming that its biosynthesis from itaconate is not restricted to immune cells (Fig. 1f). To resolve compartment-specific pathways, we permeabilized the plasma membrane that enables selective studies on itaconate metabolism in mitochondria. We traced permeabilized cells with [U-^13^C_5_]itaconate and detected fully labeled (M5) itaconate, mesaconate, citramalate, and 2HMS within mitochondria (Fig. 1g). This implicates mitochondria as the primary site of itaconate conversion pathway and synthesis of the previously overlooked compound 2HMS (Fig. 1h).

### Isotopic tracing reveals 2HMS synthesis through ACOD1 pathway in macrophages

Our ^13^C tracing studies revealed that 2HMS derives from itaconate and retains its carbon backbone indicating that probing itaconate metabolism will elucidate the 2HMS synthesis pathway in macrophages. Thus, we performed isotopic tracing experiments using [U-^13^C_6_]glucose and [U-^13^C_5_]glutamine in LPS-activated macrophages and quantified labeling on metabolites using GC-MS. Both tracers robustly labeled itaconate, mesaconate, 2HMS, and TCA cycle intermediates (Fig. 2a, b). Addition of unlabeled, ^12^C itaconate diluted labeling on itaconate, mesaconate, and 2HMS while other TCA cycle intermediates were less affected. These data further support that mesaconate and 2HMS are derived from itaconate. To gain more insights into the synthesis pathway we analyzed the labeling patterns on 2HMS from specific tracers. We observed that 2HMS, itaconate, and mesaconate all exhibited high M1 labeling from [U-^13^C_6_]glucose consistent with decarboxylation of cis-aconitate catalyzed by ACOD1 (Fig. 2c, Fig. S4a, b). Labeling on 2HMS and itaconate increased in a time-dependent manner (Fig. S4c, d). [U-^13^C_5_]glutamine tracing resulted in robust M4 labeling on these metabolites supporting synthesis from glutamine via oxidative TCA cycle metabolism (Fig. 2d, Fig. S4e, f). Since itaconate is also synthesized via the reductive carboxylation pathway, we utilized [1-^13^C_1_]glutamine which selectively labels metabolic flux through reductive carboxylation^11,35^. We observed M1 labeling on itaconate, mesaconate, and 2HMS confirming that these C_5_ dicarboxylate compounds are also synthesized through the reductive pathway (Fig. 2e).

**Figure 2:**
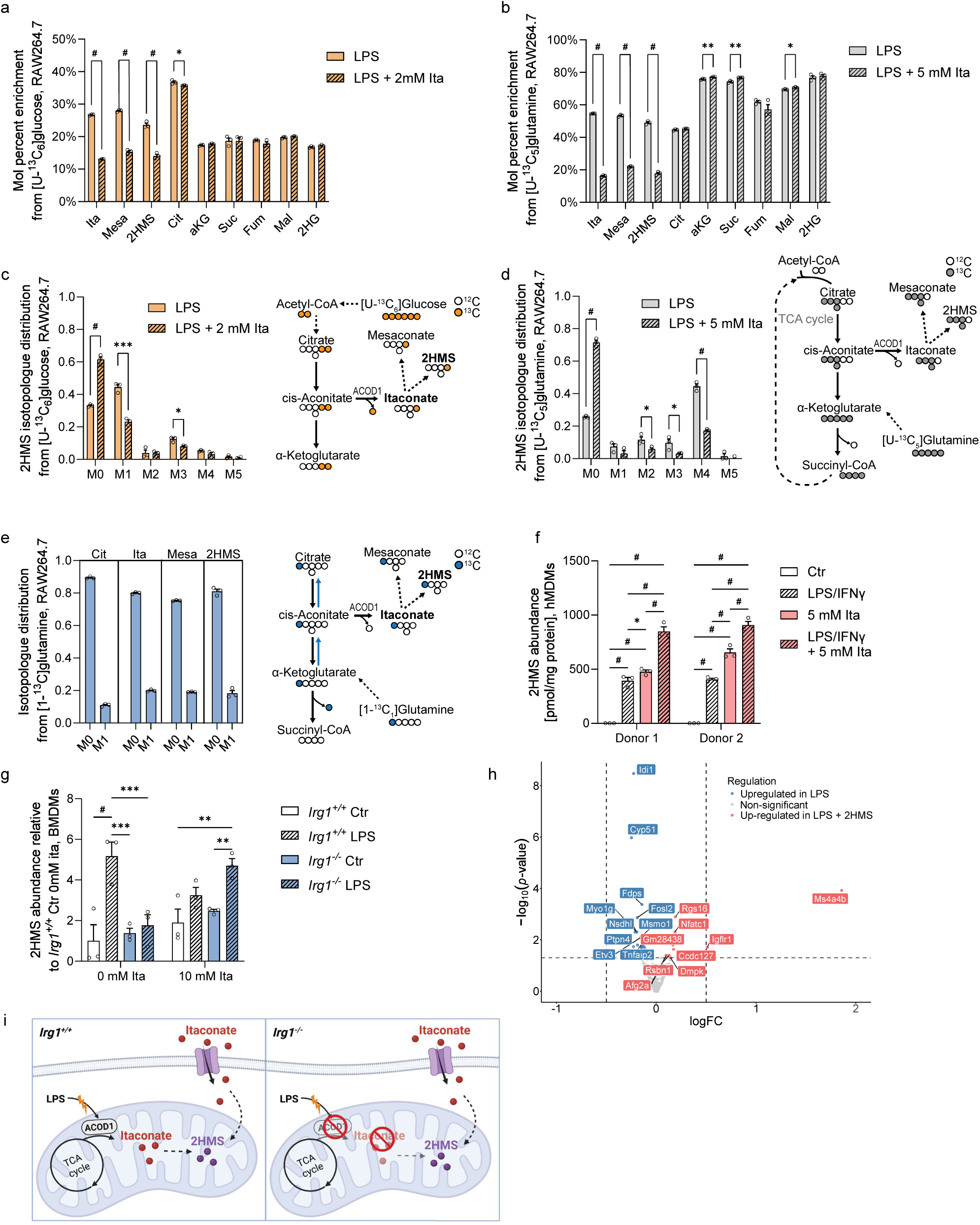
Isotopic tracing reveals 2-hydroxymethylsuccinate (2HMS) synthesis through ACOD1 pathway in macrophages. **a,** Label (mol percent enrichment) on metabolites from [U-^13^C_6_]glucose in RAW264.7 cells cultured for 24 h with 10 ng/ml LPS and 2 mM itaconate. **b,** Label (mol percent enrichment) on metabolites from [U-^13^C_5_]glutamine in RAW264.7 cells cultured for 24 h with 10 ng/ml LPS and 5 mM itaconate. **c**, 2HMS isotopologue distribution in RAW264.7 cells cultured for 24 h with [U-^13^C_6_]glucose, with metabolic map depicting atom transitions (orange cycle for ^13^C, light grey for ^12^C). **d**, 2HMS isotopologue distribution in RAW264.7 cells cultured for 24 h with [U-^13^C_5_]glutamine, with metabolic map depicting atom transitions (dark grey cycle for ^13^C, light grey for ^12^C). **e**, 2HMS isotopologue distribution in RAW264.7 cells cultured for 24 h with 10 ng/ml LPS from [1-^13^C_1_]glutamine, with metabolic map depicting atom transitions (blue cycle for ^13^C, light grey for ^12^C). **f,** 2HMS abundance in human monocyte-derived macrophages (hMDMs) cultured for 24 h with 100 ng/ml LPS, 400 U/ml interferon y (IFNy), and 5 mM itaconate. **g,** 2HMS abundance in bone marrow-derived macrophages (BMDMs) obtained from immune responsive gene 1 deficient (*Irg1*^-/-^) and wildtype (*Irg1*^+/+^) mice and cultured for 24 h with 10 ng/ml LPS and 0 mM or 10 mM itaconate. **h,** Volcano plot depicting RNA-seq results with differentially expressed genes in RAW264.7 cells cultured with 500 μM 2HMS for 4 h before stimulation with 10 ng/ml LPS for 3 h compared to control LPS conditions. **i**, Schematic depicting that wildtype (*Irg1^+/+^*) macrophages synthesize itaconate through ACOD1 activity, while *Irg1-*deficient (*Irg1^-/-^*) macrophages synthesize 2HMS from exogenous itaconate. Prepared with Biorender. Data are presented as means ± s.e.m. with three cellular replicates. Each experiment was repeated independently three times (a-d), with three donors (f), or with *n* = 3 mice (g) with similar results. *P* values were calculated by multiple unpaired *t*-test (a-d) or two-way ANOVA (f, g) with * *p* < 0.05; ** *p* < 0.01, *** *p* < 0.001, ^#^ *p* < 0.0001.

Next, we analyzed 2HMS biosynthesis in primary macrophages. We observed that human monocyte-derived macrophages (hMDMs) synthesized 2HMS, itaconate, and mesaconate in response to LPS/IFNγ stimulation and itaconate treatments further increased these levels (Fig. 2f, Fig. S4g, h). ^13^C glucose tracing in hMDMs revealed high M1 label on itaconate and 2HMS suggesting that 2HMS is derived from itaconate (Fig. S4i). We also observed similar metabolite labeling in primary bone marrow-derived macrophages (BMDMs) following LPS stimulation (Fig. S4j) confirming *de novo* 2HMS biosynthesis under inflammatory conditions through itaconate pathway.

Since itaconate synthesis is catalyzed by *Irg1*, we hypothesized that altered ACOD1 activity and *Irg1* expression may influence 2HMS synthesis. LPS-activated macrophages from *Irg1*-deficient, knockout (*Irg1*^⁻/⁻^) mice synthesized less itaconate and mesaconate that correlated with diminished 2HMS abundance compared to wildtype mice (*Irg1*^+/+^) (Fig. 2g, Fig. S4k, l). Addition of itaconate increased 2HMS levels suggesting that macrophages with impaired ACOD1 activity may rely on imported itaconate for 2HMS synthesis. We also observed that treatment with the ACOD1 inhibitor citraconate^36^ suppressed the synthesis of both itaconate and mesaconate. This inhibition reduced 2HMS levels further implicating a metabolic link between these pathways in LPS-activated macrophages (Fig. S4m). To explore the biological role of 2HMS, we treated macrophages with 2HMS and performed RNA-seq analysis. This revealed only minor changes in gene expression suggesting limited immunological effects of 2HMS in our culture system (Fig. 2h). Collectively, our findings establish 2HMS as a mammalian metabolite produced in macrophages during inflammation via the ACOD1-itaconate pathway (Fig. 2i).

### Itaconate degradation to 2HMS and mesaconate is cell-type specific

Given the lack of major transcriptional effects in macrophages, we broadened our analysis and assessed 2HMS synthesis from itaconate across cell types from different tissue origins including A549 (lung tissue-like), Huh7 (liver tissue-like), SH-SY5Y (neuronal tissue-like), RAW264.7 (macrophage-like), and 3T3-L1 (pre-adipocytes) cells. In the absence of exogenous itaconate, levels of intracellular mesaconate and 2HMS were negligible (Fig. 3a, b). Further, mesaconate and 2HMS were detectable in all cell lines tested after 24 h of culture in the presence of itaconate (Fig. 3a, b). These data confirm that 2HMS is synthesized only in the presence of itaconate and that these cells have low basal itaconate levels. We observed that diverse cell types exhibited inverse associations between mesaconate and 2HMS levels. Specifically, hepatocyte-derived Huh7 cells had high mesaconate but lower 2HMS, while 3T3-L1 cells had low mesaconate and higher 2HMS levels (Fig. 3a, b). The ratio of 2HMS to mesaconate revealed that RAW264.7 cells produce approximately similar amounts of 2HMS and mesaconate while 3T3-L1 cells synthesize higher 2HMS and Huh7 cells lower levels (Fig. 3c). We also observed robust 2HMS and mesaconate levels after 12 h of itaconate treatment (Fig. S5a, b, c). These data reveal that the enzymatic machineries for 2HMS synthesis pathway are conserved across cell types but highlight cell-specific regulation mechanisms for itaconate conversion.

**Figure 3:**
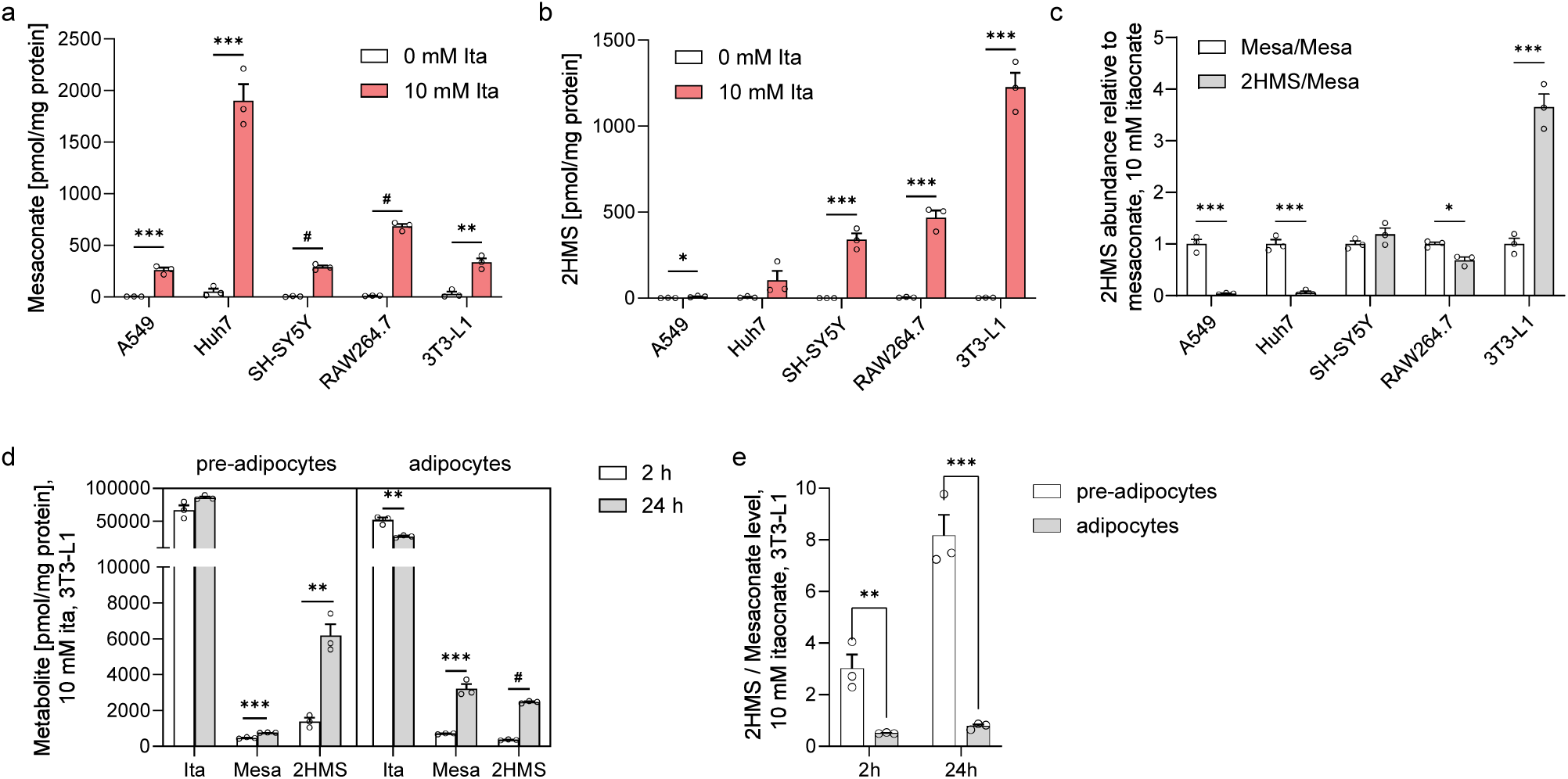
Itaconate degradation to 2-hydroxymethylsuccinate (2HMS) and mesaconate is cell-type specific. **a,** Mesaconate abundance in different cell types cultured with 0 mM or 10 mM itaconate for 24 h. **b,** 2HMS abundance in different cell types cultured with 0 mM or 10 mM itaconate for 24 h. **c,** 2HMS relative to mesaconate abundance in different cell types cultured with 10 mM itaconate for 24 h. **d,** Metabolite abundance in 3T3-L1 pre- and adipocytes cultured for 2 and 24 h with 10 mM itaconate. **e**, 2HMS relative to mesaconate abundance in 3T3-L1 pre- and adipocytes cultured for 2 and 24 h with 10 mM itaconate. Data are presented as means ± s.e.m. with three cellular replicates. Each experiment was repeated independently two times with similar results. *P* values were calculated by multiple unpaired *t*-test with * *p* < 0.05; ** *p* < 0.01, *** *p* < 0.001, ^#^ *p* < 0.0001.

Itaconate is metabolized to itaconyl-CoA which inhibits methylmalonyl-CoA mutase (MUT) activity by altering its vitamin B_12_ cofactor (cobalamin). Thus, the itaconyl-CoA–mediated MUT inhibition disrupts branched-chain amino acid (BCAA) catabolism which is highly active in adipose tissue^30,37,38^. We used adipocyte cultures with 3T3-L1 cells and compared 2HMS synthesis in differentiated cells compared to pre-adipocytes. 2HMS levels were highest in pre-adipocytes while adipocytes accumulated more mesaconate (Fig. 3d, e). Vitamin B_12_ supplementation in 3T3-L1 cultures did not alter 2HMS levels suggesting that MUT-dependent BCAA catabolism has minor effects on 2HMS pathway under these conditions (Fig. S5d).

Itaconate is biotechnologically produced by fermentation of *Aspergillus terreus* via the enzyme *cis*-aconitate decarboxylase (CAD), encoded by the *cadA* gene. To investigate potential itaconate degradation pathways, we profiled biomass and media metabolites in itaconate-producing *A. pseudoterreus* cultures and observed increasing levels of itaconate, mesaconate, and 2HMS over time (Fig. S5e, f). In contrast, these metabolites were not detectable in *A. niger* cultures due to the lack of *cadA* gene^39^. Collectively, our data suggest that itaconate degradation pathway might be evolutionarily conserved but is subject to cell-type-specific regulatory mechanisms that influence 2HMS synthesis^40^.

### 2HMS is synthesized from itaconate via a CoA - independent conversion (CIC) pathway

To identify potential enzymes involved in 2HMS synthesis pathway, we targeted methylglutaconyl-CoA hydratase (AUH) encoded by methylglutaconyl-CoA hydratase (*Auh*), and succinyl-CoA synthetase (SCS) encoded by succinate-CoA ligase (*Suclg1*) in 3T3-L1 cells which synthesize high amounts of 2HMS (Fig. 4a, Fig. S6a, b). Both enzymes have been linked to the itaconate degradation pathway, specifically for mesaconate synthesis^29,31,32,41^, but whether they are involved in 2HMS synthesis is not known. We observed increased itaconate and decreased mesaconate levels upon silencing of *Suclg1* indicating impaired mesaconate synthesis via itaconyl-CoA-dependent pathway. The 2HMS levels were also increased upon *Suclg1* silencing suggesting that itaconate might be redirected for conversion to 2HMS when SCS activity is impaired. In contrast, impaired activity of AUH decreased 2HMS levels indicating that AUH activity is required for 2HMS synthesis. We also observed increased methylmalonate levels in response to itaconate treatment due to impaired MUT activity in the presence of itaconate (Fig. 4b). Impaired AUH activity further increased levels of methylmalonate suggesting increased flux through CoA-dependent conversion (CDC) pathway. In addition to SCS, succinyl-CoA glutarate CoA-transferase (SUGCT) has also been described to catalyze the synthesis of itaconyl-CoA from itaconate as an alternative pathway^42^. Thus, we targeted *Sugct* expression and observed slightly increased 2HMS abundances while mesaconate levels were unaffected. These data indicate minor involvement of SUGCT activity in itaconate degradation pathway in our culture model (Fig. S6c, d).

**Figure 4:**
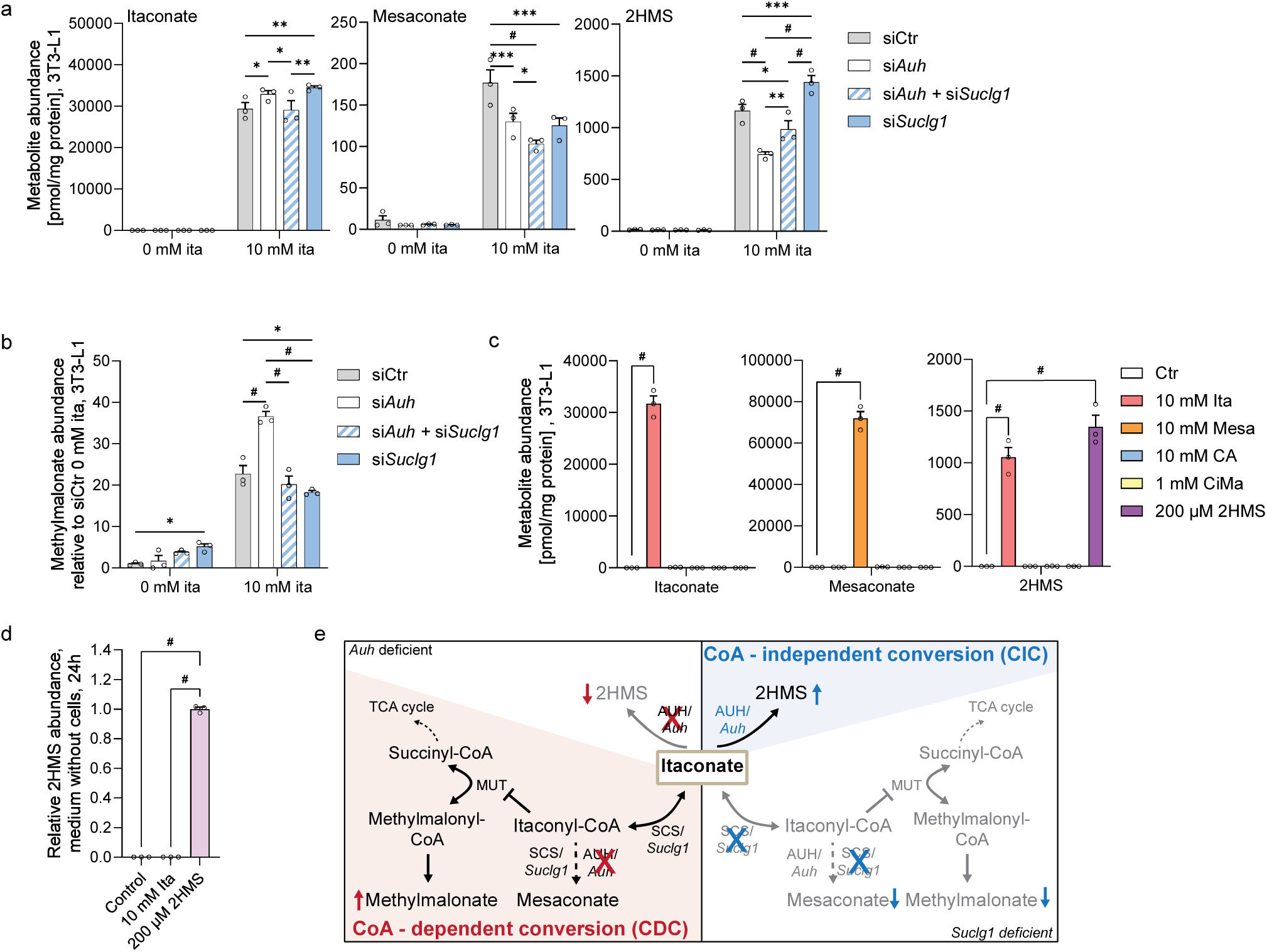
2-hydroxymethylsuccinate (2HMS) is synthesized from itaconate via an itaconyl-CoA independent conversion (CIC) pathway. **a,** Metabolite abundance in 3T3-L1 cells with small interfering RNA (siRNA)-mediated gene knockdown targeting methylglutaconyl-CoA hydratase (si*Auh*), succinate-CoA ligase (si*Suclg1*), or both genes combined (si*Auh* + si*Suclg1*) compared to control condition (siCtr) cultured with 0 mM or 10 mM itaconate for 24h. **b,** Methylmalonate levels in 3T3-L1 cells in response to siRNA-mediated gene knockdown of *Auh* and *Suclg1* compared to control conditions (siCtr) cultured with 0 mM or 10 mM itaconate for 24h. **c,** Metabolite abundance in 3T3L-1 cells cultured for 24h with itaconate (ita), mesaconate (mesa), citraconate (CA), citramalate (CiMa) or 2-hydroxymethylsuccinate (2HMS) compared to untreated control cells (Ctr). **d,** 2HMS abundance in cell-free complete DMEM medium after 24 h incubation with 10 mM itaconate or 200 µM 2HMS. **e,** Schematic depicting succinyl CoA synthetase (SCS) mediated CoA-dependent conversion (CDC) pathway, methylglutaconyl-CoA hydratase (AUH) mediated CoA-independent conversion (CIC) pathway, and impact on methylmalonyl-CoA mutase (MUT) activity. Data are presented as means ± s.e.m. with three cellular replicates. Each experiment was repeated independently three (a, b) or two (c, d) times with similar results. *P* values were calculated by two-way ANOVA (a, b) or one-way ANOVA compared to Ctr condition (c, d) with * *p* < 0.05; ** *p* < 0.01, *** *p* < 0.001, ^#^ *p* < 0.0001.

Further, we excluded potential synthesis of 2HMS from other C_5_ dicarboxylate compounds including mesaconate, citramalate, and citraconate (Fig. 4c). These data demonstrate that 2HMS is not an intermediate in the mesaconate or citramalate pathways. The synthesis of 2HMS and mesaconate appears to be unidirectional implying that these metabolites are not interconverted. We also observed similar effects in RAW264.7 macrophage-like cells, confirming that 2HMS is synthesized from itaconate through an AUH-dependent pathway (Fig. S6e-h). Collectively, our pathway analysis reveals that 2HMS biosynthesis occurs via a CoA-independent conversion (CIC) pathway catalyzed by AUH. This CIC pathway is distinct from the CoA-dependent conversion (CDC) pathway which requires SCS for itaconyl-CoA and mesaconate synthesis (Fig. 4e).

### CpG-induced inflammation drives the synthesis of itaconate and 2HMS in vivo

As we observed the apparent production of 2HMS from itaconate *in vitro*, we also wondered whether this pathway would be active *in vivo*. Given the close relationship between itaconate and macrophage activation, we employed a mouse model of macrophage activation syndrome to stimulate itaconate production using the CpG oligodeoxynucleotide ODN1826, an agonist of Toll-Like Receptor 9 (TLR9)^43–45^. We observed a strong increase in plasma itaconate and mesaconate levels following CpG treatment while 2HMS levels remained unchanged (Fig. 5b). These data suggest that TLR9 stimulus triggers inflammation-dependent synthesis of itaconate *in vivo*.

**Figure 5:**
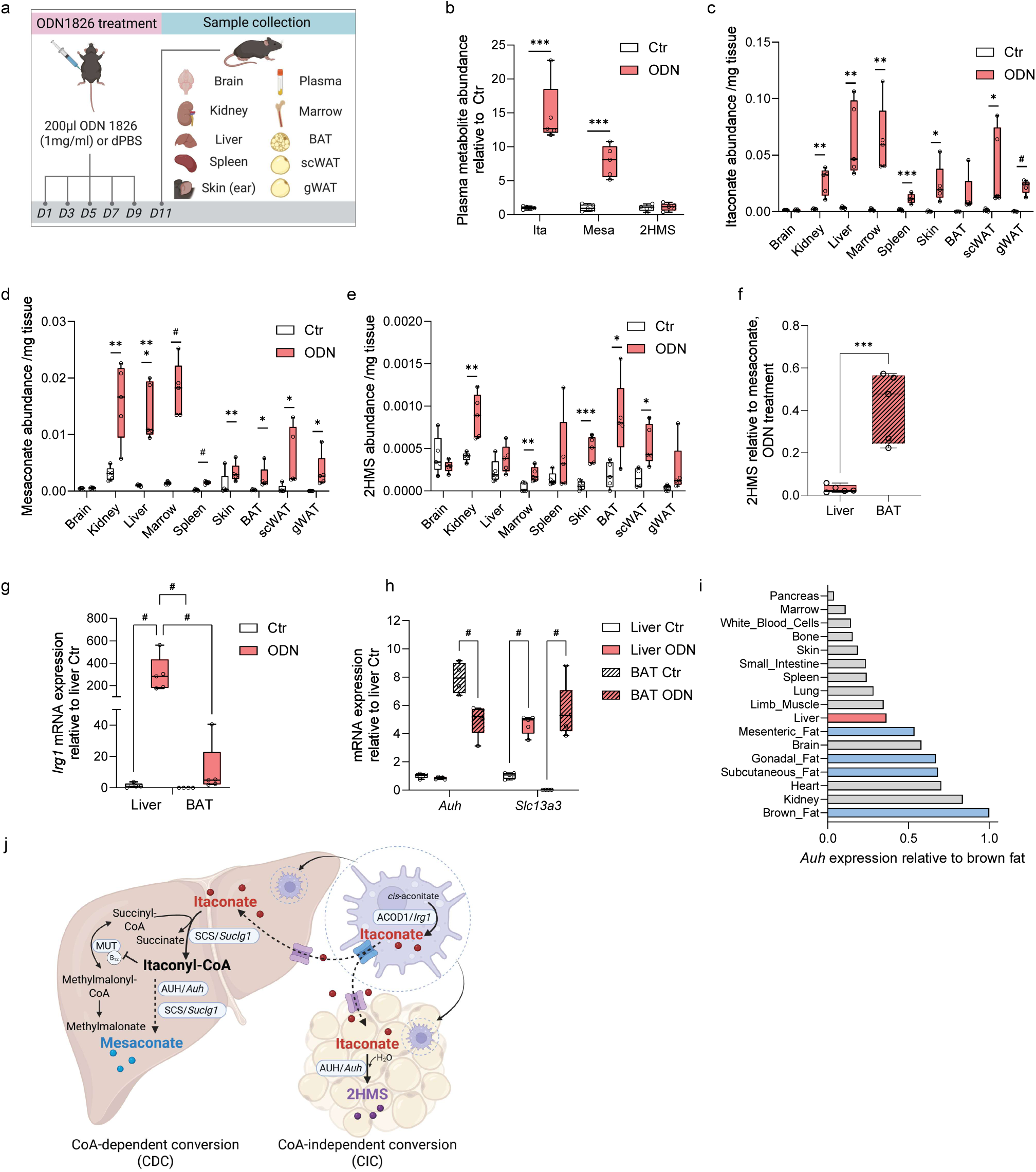
Itaconate and 2-hydroxymethylsuccinate (2HMS) are synthesized *in vivo* in an animal model of CpG oligodeoxynucleotide (ODN) induced inflammation. **a,** Schematic depicting experimental setup for CpG ODN1826-induced inflammation. **b,** Itaconate, mesaconate, and 2HMS abundances in plasma. **c,** Itaconate abundance in tissues **d,** Mesaconate abundance in tissues. **e**, 2HMS abundance in tissues. **f**, 2HMS relative to mesaconate levels in CpG-treated liver and BAT tissues g, *Irg1* expression level in liver and BAT tissue. **h**, *Auh* and *Slc13a3* expression level in liver and BAT tissue. **i**, *Auh* expression in various tissues (Schaum et al., Nature, 2020). **j**, Schematic depicting itaconate degradation pathways in liver and adipose tissue with distinct pathways for mesaconate and 2HMS synthesis. Data are presented as box (25^th^ to 75^th^ percentile with median line) and whiskers (min. to max. values). Animals were injected with 200 µg CpG oligonucleotide every second day for a total of 5 injections (ODN, *n* = 5) and compared to control conditions (Ctr, *n* = 5). Schematics in Figures 5a and 5j were prepared with Biorender. BAT, brown adipose tissue; scWAT, subcutaneous white adipose tissue; gWAT, gonadal white adipose tissue. *P* values were calculated by multiple unpaired *t*-test (b-e), unpaired *t*-test (f), one-way ANOVA (g), or two-way ANOVA compared to Ctr (h) with * *p* < 0.05; ** *p* < 0.01, *** *p* < 0.001, ^#^ *p* < 0.0001.

Since macrophages are distributed in diverse tissues and 2HMS is synthesized from itaconate in diverse cell types *in vitro* (Fig. 3b, Fig. S5c), we also measured these metabolite levels across tissues. We observed that itaconate accumulated significantly in kidney, liver, and bone marrow. Itaconate levels in adipose tissue (AT), including brown AT (BAT), subcutaneous white AT (scWAT), and gonadal white AT (gWAT), were lower compared to liver tissue (Fig. 5c). Further, we detected high mesaconate levels in kidney, liver, and bone marrow that correlated with itaconate levels in these tissues (Fig. 5d). Notably, 2HMS levels were significantly increased in BAT as well as kidney, bone marrow, skin, and scWAT but not in liver after CpG treatment (Fig. 5e). These data suggest divergent, tissue-specific conversion pathways of itaconate yielding mesaconate and 2HMS, specifically between liver and BAT tissue (Fig. 5f).

Since we detected high itaconate levels in liver tissue following CpG treatment we examined expression of *Irg1* which encodes the itaconate-synthesizing enzyme ACOD1. Treatment significantly increased hepatic *Irg1* expression while expression in BAT tissue was less affected suggesting liver myeloid cells drive itaconate production in our CpG-induced inflammation model (Fig. 5g). The low induction of *Irg1* in BAT implies potential itaconate uptake mechanisms in this tissue. We therefore quantified expression of *Slc13a3* encoding sodium-dependent dicarboxylate transporter 3 (SLC13A3, NaDC3) which facilitates the transport of itaconate^46–48^. We observed higher base levels of *Slc13a3* in liver compared to BAT. *Slc13a3* expression significantly increased in response to CpG treatment in both, BAT and liver tissue (Fig. 5h). These data indicate that inflammation may enhance itaconate uptake into tissues with less endogenous, ACOD1-driven itaconate synthesis. We also observed significantly increased *Auh* levels in BAT compared to liver tissue that may promote 2HMS synthesis in a CoA-independent manner (Fig. 5h). Finally, we reanalyzed published RNA-seq data^49^ and observed that *Auh* expression was highest in adipose tissues (Fig. 5i). In contrast, the liver favored the CoA-dependent conversion (CDC) pathway converting itaconate to mesaconate further indicating distinct pathways for itaconate conversion (Fig. 5j). This CDC route is linked to B_12_ metabolism through itaconyl-CoA suggesting a potential link to hepatic B_12_ homeostasis. Our findings demonstrate that tissues employ distinct mechanisms to metabolize itaconate in a tissue-specific manner (Fig. 5j).

### 2HMS is synthesized from itaconate in vivo

In our previous study, we traced the fate of [U-^13^C_5_]itaconate *in vivo* and demonstrated rapid renal clearance of plasma itaconate, its conversion to mesaconate and citramalate, and partial fueling into the TCA cycle metabolism in liver and kidney^28^. The ^13^C itaconate tracing revealed the highest mesaconate levels in the kidneys suggesting tissue-specific metabolism^28^. To determine whether 2HMS is synthesized from itaconate *in vivo*, we reanalyzed our data and quantified labeling on 2HMS. We observed labeled 2HMS in the plasma upon tracer administration indicating *de novo* synthesis of 2HMS from [U-^13^C_5_]itaconate (Fig. 6a). Plasma 2HMS peaked 15 min post-injection and declined within 45 minutes mirroring the kinetics of ^13^C labeled itaconate and mesaconate (Fig. 6a). Labeled 2HMS also accumulated in kidney, liver, lung, brain, and spleen. We detected the highest 2HMS levels in kidney and liver at 15 min that correlated with tissue itaconate and mesaconate abundances (Fig. 6b). Urinary 2HMS was predominantly M5 labeled from [U-^13^C_5_]itaconate (Fig. 6c,d). Notably, since unlabeled 2HMS was undetectable in the urine, systemic 2HMS pools mainly originated from itaconate and were cleared by the kidneys. Collectively, our data identified 2HMS as a previously unrecognized metabolite synthesized from itaconate *in vivo*. Since systemic abundances of 2HMS and itaconate are dynamically influenced by renal excretion, levels of C_5_ dicarboxylate compounds might be regulated in disease states.

**Figure 6:**
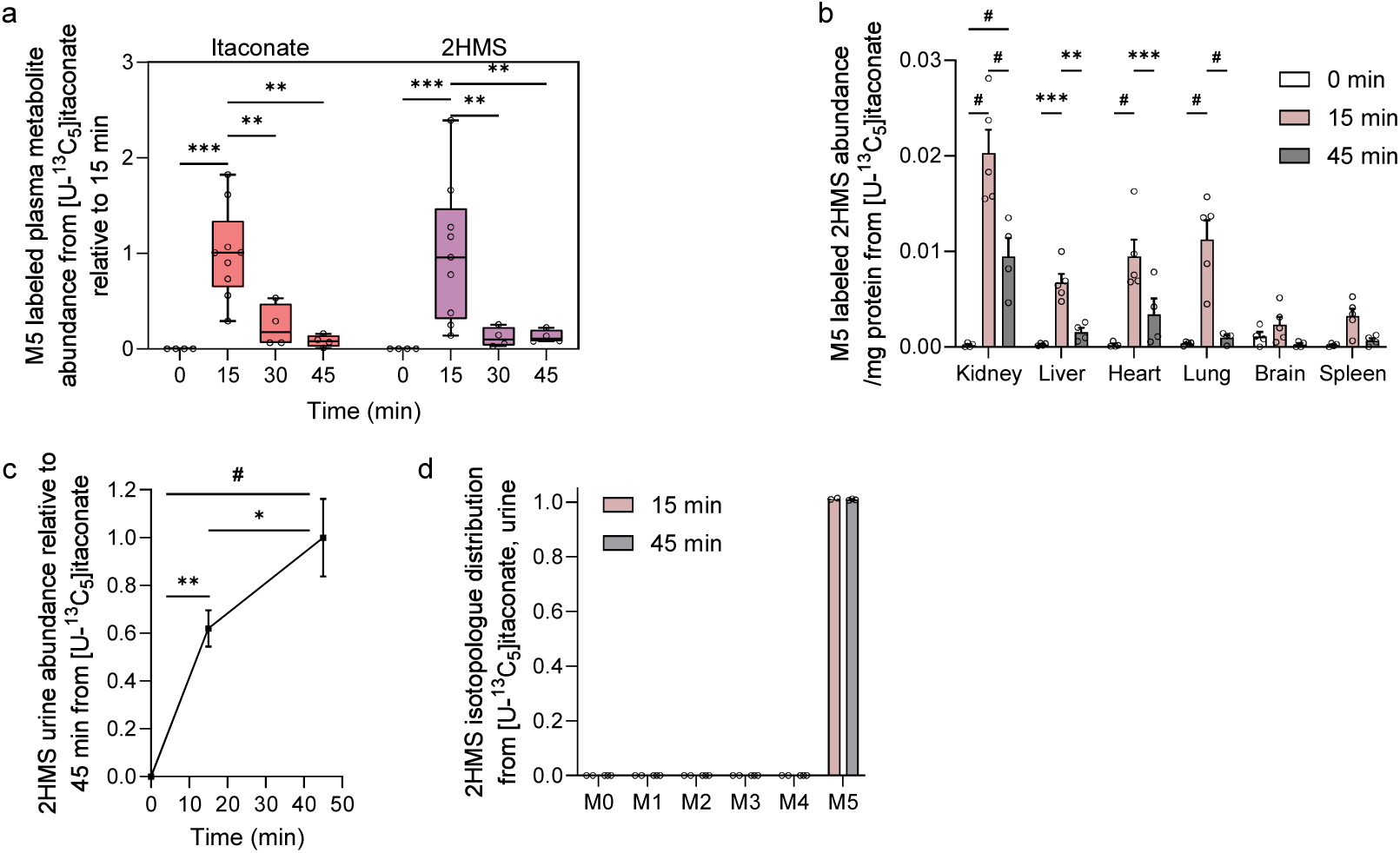
^13^C itaconate is metabolized to 2-hydroxymethylsuccinate (2HMS) and further cleared renally *in vivo*. **a,** Labeled plasma metabolite abundance. **b,** Labeled metabolite abundance in tissues. **c,** Labeled 2HMS abundance in urine. **d,** Urine 2HMS isotopologue distribution. Mice were injected with 400 mg/kg body weight [U-^13^C_5_]itaconate. Data are presented as box (25^th^ to 75^th^ percentile with median line) and whiskers (min. to max. values) (a) or means ± s.e.m. (b-d). Plasma samples at 0 min (n = 4), 15 min (n = 9), 30 min (n = 4), and 45 min (n = 4); tissue samples at 0 min (n = 4), 15 min (n = 5), 45 min (n = 4), urine samples at 0 min (n = 3), 15 min (n = 2), and 45 min (n = 3). *P* values were calculated by two-way ANOVA (a,b) and compared to 0 min itaconate (c) with * *p* < 0.05; ** *p* < 0.01, *** *p* < 0.001, ^#^ *p* < 0.0001.

## Discussion

Here, we identify 2HMS as a previously overlooked mammalian metabolite that is synthesized *de novo* from itaconate under inflammatory conditions in macrophages. 2HMS synthesis occurred in mitochondria via an AUH-dependent, CoA-independent conversion pathway that is distinct from the CoA-dependent mesaconate pathway. These tissue-specific pathways of itaconate degradation suggest evolutionary adaptation to balance immune responses. The separate enzymatic machineries for mesaconate and 2HMS synthesis demonstrate a metabolic bifurcation with implications for TCA cycle, amino acid, and vitamin B_12_ metabolism.

One study postulated the existence of 2HMS (referred to as itamalate) in the 1950s and limited subsequent research on 2HMS is availabe^31^. Zimmermann et al. have recently detected 2HMS in a mixture after generating it from itaconate using an iron catalyst, though the compound was not assigned a specific name in their study^50^. One reason it has been overlooked might be due to the structural similarity and identical masses to other C_5_ dicarboxylates including 2HG, 3HG, 3-methylmalate, and citramalate. These compounds are difficult to differentiate with automated mass spectrometry methods and require sensitive instrumentation for low abundant metabolites. Our analytical methods distinguished 2HMS from structurally related compounds and confirmed 2HMS as a distinct itaconate-derived metabolite that is synthesized during inflammatory conditions in macrophages. We also synthesized an authentic 2HMS standard to validate its identity. Our discovery may open new research directions and mitigate potential misidentification of C_5_ dicarboxylates in future studies.

Our results and permeabilized tracing demonstrate that the conversion of itaconate to 2HMS proceeds via a CoA-independent pathway in mitochondria. This pathway is particularly active in adipose tissue with high 2HMS levels, whereas the liver predominantly synthesizes mesaconate through a CoA-dependent route. These tissue-specific differences are supported by differential expression of key enzymes as well as our *in vitro* studies in hepatocyte cultures. BAT expresses high levels of both AUH and the itaconate transporter SLC13A3 to uptake circulating itaconate and to subsequently convert it to 2HMS. The liver favors the CoA-dependent conversion (CDC) pathway for itaconyl-CoA formation catalyzed by SCS (Fig. 5j). The transfer of CoA from succinyl-CoA to itaconyl-CoA intersects with substrate-level phosphorylation and TCA cycle metabolism that may further affect mitochondrial function^29,51^. Of note, 2HMS synthesis bypasses the synthesis of itaconyl-CoA which inactivates the B_12_-dependent enzyme MUT involved in BCAA catabolism. Thus, 2HMS synthesis and bypassing itaconyl-CoA formation may preserve TCA cycle metabolism and B_12_-dependent metabolic processes. This adaptation may also influence adipose function and BAT thermogenesis by maintaining metabolic fluxes^52^. Consequently, the balance between these two itaconate conversion (CIC and CDC) pathways may influence physiological processes during itaconate treatments or inflammatory-driven itaconate synthesis.

AUH is involved in leucine metabolism and mutations in the *Auh* gene cause the rare metabolic disease 3-methylglutaconyl-CoA hydratase deficiency (3-methylglutaconic aciduria type 1, MGCA1) with neurological symptoms^53^. Our discovery that AUH mediates 2HMS synthesis suggests a previously unrecognized metabolic axis between itaconate and BCAA metabolism that may intersect with the 2HMS synthesis pathway. The itaconate conversion product itaconyl-CoA influences BCAA and B_12_ metabolism which may contribute to pathophysiology in tissues with high AUH expression. AUH binds to AU-rich elements (AREs) to modulate post-transcriptional regulation of mRNAs^54,55,57^. Consequently, as *Irg1* contains AREs, alteration of AUH may influence *Irg1* expression, itaconate production, and biological responses^56^.

Itaconate has emerged as a therapeutic candidate for the treatment of reperfusion injuries^16,58^, inflammation^59^, cancer^1,47^, and liver diseases^17,19,52^. Our recent ^13^C itaconate *in vivo* study revealed rapid renal clearance of itaconate while a minor fraction also fueled carbons for TCA cycle metabolism^28^. However, we still do not fully understand how itaconate and its breakdown products influence cellular processes. Mammalian itaconate degradation may also influence host-pathogen interactions as certain pathogenic bacteria catabolize itaconate to reduce its antimicrobial efficacy during infection^60,61^. Future work should include investigating how 2HMS is distributed across tissues in healthy and pathogenic conditions and clarifying its effects on cellular processes.

In summary, we identify 2HMS as a distinct metabolite synthesized *de novo* from itaconate mediated by AUH activity in mitochondria. Our findings uncover the fate of itaconate and its conversion pathways and establish analytical frameworks for characterizing C_5_ dicarboxylate compounds. We further demonstrate that tissue-specific regulation of 2HMS reflects distinct mechanisms of itaconate degradation routes which may influence metabolic pathways and immune regulation. This work highlights 2HMS and itaconate metabolism as potential targets for therapeutic interventions.

## Material and methods

### Cell lines

The following cell lines were used for the experiments: RAW264.7 (TIB-71, ATCC, mouse macrophage-like cells), A549 (CCL-185, ATCC, human lung carcinoma cells), 3T3-L1 (CL-173, ATCC, mouse pre-adipocytes), HEK-293 (CRL-1573, ATCC, human embryonic kidney cells), Huh7 (kindly provided by the laboratory of Christian Metallo, Salk Institute of Biological Studies, USA, human hepatocellular carcinoma cells), HepG2 (HB-8065, ATCC, human hepatocellular carcinoma cells), and SH-SY5Y (CRL-2266, ATCC, human neuroblastoma cells). Cells were cultured in high glucose Dulbecco’s Modified Eagle Medium (DMEM, Cat.# 41965-039, Gibco), containing 25 mM glucose, 4 mM glutamine, and supplemented with 50 U/ml penicillin, 50 µg/ml streptomycin (P/S, Cat.# 15140-122, Gibco) and 10 % (v/v) Fetal Bovine Serum (FBS, FBS.SAM.0500, Bio&Cel, sourced from South America) as growth medium in a humidified cell culture incubator (Thermo Scientific) at 37 °C and 5 % CO_2_. A549, 3T3-L1, Huh7, HepG2, HEK2-293, and SH-SY5Y cells were detached using 0.05 % trypsin-EDTA (Cat.# 25300-054, Gibco). All media were adjusted to pH 7.3. Cells were tested negative for mycoplasma using MycoAlert^®^ Mycoplasma Detection Kit (Cat.# LT07-118, Lonza).

### 3T3-L1 cultures and differentiation

Murine 3T3-L1 pre-adipocytes were cultured in high glucose DMEM with 10 % FBS and cell densities were maintained below 40 % confluency. For differentiation, 40,000 cells/well were seeded into 12-well tissue culture plates (Greiner Bio-One) on day –5 and allowed to reach confluence (termed day –2). On day 0 differentiation was induced by treating cells with a differentiation cocktail containing 0.5 mM 3-isobutyl-1-methylxanthine (IBMX, Cat.# I5879, Sigma-Aldrich), 10 μM dexamethasone (Cat.# D4902, Sigma-Aldrich), and 5 μg/ml insulin (Cat.# I6634, Sigma-Aldrich) in a complete DMEM medium containing 10 % FBS. Media was replaced on day 3 with a complete DMEM medium containing 5 μg/ml insulin. From day 6 onward, cells were resting in complete medium without additional differentiation supplements. Cobalamin (Vitamin B_12_, 1 µM, Cat.# 38310, Serva) were supplemented to cultures with treatment of 10 mM itaconate for 24 h. Metabolite extractions were conducted between differentiation days 6-8 unless otherwise stated and quantified using GC-MS technology.

### Cell treatments

Cells were seeded at a density of 300,000 cells/ml in 12-well tissue plates (Greiner Bio-One) and cultured overnight to achieve a confluent monolayer. Cells were treated with itaconate (stock solution 1 M, I29204-100G, Sigma-Aldrich), mesaconate (stock solution 1 M, 131040-50G, Sigma-Aldrich), citraconate (stock solution 1 M, C82604-100G, Sigma-Aldrich), citramalate (stock solution 100 mM, 27455-1G, Sigma-Aldrich), or 3-methylmalate (stock solution 100 mM, Molport-001-785-695, molport) with concentrations and durations indicated in the respective figure legends. To induce immune responses in macrophages, RAW264.7 cells and mouse bone marrow-derived macrophages (BMDMs) were treated with 10 ng/ml lipopolysaccharide (LPS, stock 1 mg/ml, E. coli O128:B12, L2755, Sigma-Aldrich), and human monocyte-derived macrophages (hMDMs) received 100 ng/ml LPS in combination with 400 U/ml interferon-gamma (IFN-γ, stock 2,000,000 U/ml, AF-300-02-100, Peprotech). For standard addition experiments, metabolites (25 µl itaconate with a concentration of 10 mM, 10 µl 2HMS with a concentration of 1 mM) were spiked into the aqueous methanol/water fraction post metabolite extraction and prior to drying samples for GC-MS analysis.

### Bone marrow-derived macrophage (BMDM) cultures

The *Irg1*^-/-^ (also known as *Acod1*^-/-^) knockout (KO) mice were originally generated by Dr. Haruhiko Koseki at the RIKEN Institute using stem cells obtained from the Knockout Mouse Project (KOMP) Repository (Irg1tm1a(KOMP)Wtsi) and were bred at Helmholtz Centre for Infection Research (HZI). C57BL/6J mice, acquired from Charles River, were used as wild-type controls. All experiments used 6- to 12-week-old male mice. BMDMs were isolated from femurs and tibias. High glucose DMEM (Cat.# 41965-039, Gibco) was supplemented with 10 % (v/v) FBS, 50 U/ml penicillin, and 50 µg/ml streptomycin and used as growth medium. Femurs and tibias were harvested from both hind legs of each mouse, placed in ice-cold PBS, and cleaned of muscles. For marrow collection, the ends of each bone were cut, and the bones were positioned open-end down in a pipette tip in a 1.5 ml tube (containing 200 µl growth medium). Bone marrow was extruded by centrifugation at 5,200 × *g* for 30 sec. Bone marrow was gently resuspended, transferred to 15 ml reaction tubes containing 10 ml growth medium, and centrifuged at 200 x *g* for 10 min at 4 °C. Pellets were resuspended with 36 ml growth DMEM supplement with 40 ng/ml recombinant rM-CSF (premium grade, Cat.# 130-101-704, source E.coli, Miltenyi Biotec) and seeded in three 10 cm culture dishes (Cat.# 82.1473.001, Sarstedt). Medium was refreshed at day 3 with growth DMEM supplemented with 25 ng/ml rM-CSF. On day 6 or 7, differentiated BMDMs were collected and replated into 12-well plates (Greiner Bio-One) at a density of 1 million cells/ml. Cells were treated with 10 ng/ml LPS for 24 h. Metabolites were extracted on day 7 or 8 and quantified using GC-MS technology.

### Primary human peripheral blood mononuclear cell (PBMC) cultures

Primary human peripheral blood mononuclear cells (PBMCs) were differentiated from monocytes and isolated from whole human blood taken from random donors provided by the donation service at the Klinikum Braunschweig. Blood was provided in leukoreduction system (LRS) chambers. For isolation of PBMC, whole blood was rinsed out of LRS chamber and diluted in 1x PBS. For initial separation, blood/PBS suspension was stacked on 4 ml Biocoll® separation solution (density 1.077 g/ml, BS.L6115, Bio&SELL) to perform density gradient centrifugation at 700 *x g* without a break. The white PBMC ring was taken off and washed twice with PBS. For further separation, the cells were magnetically labeled with CD14 microbeads (Cat.# 130-050-201, Miltenyi Biotec) with a 1:5 ratio of cells to CD14 microbeads in MACS® buffer (autoMACS® Rinsing Solution, Cat.# 130-091-222, + 5 % MACS® BSA stock solution, Cat.# 130-091-376, Miltenyi Biotec). Cell suspension with beads was incubated for 15 min on ice before adding to theMACS® buffer equilibrated LS-Columns (Cat.# 130-042-401, Milteny Biotec) attached to a magnetic MACS® separator. Columns were rinsed 3 times with 3 ml MACS® buffer. The CD14-labeled monocytes were eluted with 3 ml MACS® buffer using a plunger.

For differentiation, monocytes were resuspended in RPMI containing 10 % FBS, 50 U/ml penicillin, 50 µg/ml streptomycin, and 50 U/ml human macrophage colony-stimulating factor (rh M-CSF, Cat.# 11343115, ImmunoTool) and cultured for 4 days in 6-well tissue-culture plates with 2*10^6^ cells/ml in a humidified cell culture incubator (Thermo Scientific) at 37 °C and 5 % CO2. Cells were treated with 100 ng/ml LPS and 400 U/ml IFNγ and 5 mM itaconate for 24 h before metabolite extraction. Samples were analyzed on GC-MS platform.

### Itaconate production in *Aspergillus*

*Aspergillus pseudoterreus* (DSM 826, Leibnitz Institute DSMZ, Germany) and *Aspergillus niger* (CBS 101705, Westerdijk Fungal Biodiversity Institute, Netherlands) were pre-cultured by inoculating spore suspensions (10^6^ spores per ml) into 20 ml pre-culture medium in 100 ml Erlenmeyer flasks. The medium contained (per L): 25 g glucose, 4.5 g MgSO4·7H₂O, 0.4 g NaCl, 4 mg ZnSO_4_·7H_2_O, 100 mg KH_2_PO_4_, 2 g NH_4_NO_3,_ and 0.5 g corn steep solids (CSS). Cultures were incubated at 30°C with agitation at 170 rpm for 3 days. On day 0 (D0), the full 20 ml pre-culture was transferred into 200 ml of CAD production medium in 1 liter Erlenmeyer flasks. The CAD medium was modified from the formulation described by von der Straat et al. and contained (per liter): 25 g glucose, 0.5 g CSS, 3.45 g MgCl_2_, 1.5 g MgSO_4_·7H_2_O, 3 g NH_4_NO_3_, 0.4 g NaCl, 33 mg ZnSO₄, 50 mg KH_2_PO_4_, 1 g CaCl_2_, and 60 mg CuSO₄. The pH was adjusted to 2.0 before inoculation. Fermentations were carried out at 30°C with shaking at 170 rpm. From D0 to D4, 1 ml suspension culture was collected daily, centrifuged at 13,000 *x* g for 2 min, and the clarified supernatants and the biomass were stored at −20°C for subsequent metabolite extraction.

### Isotopic tracing analysis

Tracing of cell lines and BMDMs was performed in DMEM (Cat.# D5030, Sigma-Aldrich) supplemented with 3.7 g/l bicarbonate. For ^13^C glucose tracing experiments, unlabeled glucose was replaced with 25 mM [U-^13^C_6_]glucose (Cat.# CLM-1396-1, Cambridge Isotope Laboratories). For ^13^C glutamine tracing experiments, unlabeled glutamine was replaced with either 4 mM [U-^13^C_5_]glutamine (Cat.# CLM-1822-H-0.5, Cambridge Isotope Laboratories) or [1-^13^C_1_]glutamine (Cat.# CLM-3612-1, Cambridge Isotope Laboratories). For tracing experiments with hMDMs, cells were cultured in SILAC RPMI 1640 Flex Media (Cat.# A24942-01, Gibco) containing 11.1 mM [U-^13^C_6_]glucose and 2 mM unlabeled glutamine. All tracer media were sterile-filtered through 0.22 µm filters and supplemented with 10 % FBS, 50 U/ml penicillin, and 50 µg/ml streptomycin. For inflammatory stimulation, cells were stimulated with LPS as indicated in each figure legend. ^13^C itaconate tracing studies were performed by supplementing [U-^13^C_5_]itaconate (NIH Common Fund Metabolite Standards Synthesis Core, NHLBI Contract No. HHSN268201300022C) to the cultures as indicated in each figure legend. Labeling on metabolites from ^13^C tracer was quantified using GC-MS platform.

### Permeabilized cell trace

Cells were seeded on a 12-well culture plate and permeabilized with recombinant perfringolysin O (rPFO, commercially XF Plasma membrane permeabilizer (PMP), Agilent Technologies, Cat.# 102504-100) to allow access to mitochondrial metabolism. The mitochondrial assay medium contained 125 mM sucrose, 65 mM KCl, 5 mM KH_2_PO_4_, 20 mM HEPES, 1 mM MgCl_2_, 0.5 mM EGTA, and 0.2 % BSA with pH adjusted to 7.3. Culture media was removed and cells were washed with PBS. Cells were then rinsed twice with 0.2 ml mitochondrial assay medium. Next, cells were traced for 20 min in a non-CO_2_ incubator at 37 °C in mitochondrial assay medium supplemented with 1 mM glutamine, 1 mM malate, 4 mM ADP, and 1 or 5 mM [U-^13^C_5_]itaconate. After incubation, cells were then washed twice with 150 mM NaCl and used for metabolite extraction following the protocol described above for subsequent GC-MS analysis.

### Metabolite extraction for GC-MS analysis

Metabolites were extracted as previously described in detail^62^. Briefly, cells were washed with saline solution (0.9 % w/v NaCl) and quenched with 0.25 ml −20 °C methanol. After adding 0.1 ml 4°C cold water containing an internal standard norvaline (5 µg/ml). The mixture was collected into a pre-cooled microcentrifuge tube, and 20 µl of the methanol-water mixture was transferred into a 96-well plate for protein quantification. Next, 0.25 ml −20 °C chloroform was added to the sample, and the extracts were vortexed for 10 min at 4 °C and centrifuged at 16,000 × *g* for 5 min at 4°C. The upper aqueous phase was evaporated under vacuum at 4 °C and used for GC-MS analysis. Medium samples were centrifuged at 300 x *g* for 5 min at 4°C, then 10 µl of medium was mixed with 90 µl of extraction fluid (methanol:water is 9:1, containing 4.44 µg/ml norvaline), vortexed for 15 sec, and spun again at 17,000 x *g* for 5 min at 4°C. 60 µl of the supernatant was dried under vacuum at 4°C and used for GC-MS analysis.

For the CpG oligonucleotides (ODN1826) inflammation-induced *in vivo* model, plasma metabolite levels were extracted using 20 µl plasma and 90 µl methanol:water (9:1) containing 5 µg/ml norvaline as internal standard. Tissues obtained from the CpG *in vivo* model were pulverized using a cell crusher cryogenic tissue pulverizer (Cellcrusher, Cork, Ireland). For tissue extraction, 10 - 20 mg pulverized tissue was homogenized with a ball mill (Retsch Mixer Mill MM 400) at 30 Hz for 2 min and metabolites were extracted with 0.5 ml −20 °C methanol, 0.2 ml 4°C cold water (containing 5 µg/ml internal standard norvaline), and 0.5 ml −20 °C chloroform. The plasma and tissue extracts were vortexed for 10 min at 4 °C and centrifuged at 17,000 × *g* for 5 min at 4 °C. The upper aqueous phase was evaporated under vacuum at 4 °C and used for GC-MS analysis.

For *Aspergillus* biomass extraction, 10 - 20 mg of fungal biomass was mixed with 0.25 ml −20°C methanol and 0.1 ml water (containing 5 µg/ml norvaline), homogenized by a ball mill (30 Hz, 30 sec), and 0.25 ml −20°C chloroform was added. The sample was vortexed for 10 min at 4°C. 10 µl *Aspergillus* culture media was used for metabolite extraction and mixed with 90 µl cold extraction fluid (methanol:water 9:1 with 4.44 µg/ml norvaline), vortexed for 15 s. Both sample types were centrifuged at 17,000 x *g* for 5 min at 4°C, the upper aqueous phase was dried under vacuum at 4°C, and used for GC-MS analysis.

### Gas Chromatograph–Mass Spectrometry (GC-MS) analysis and sample preparation

Metabolites were analyzed and quantified, as previously described in detail^62^. Polar metabolites were derivatized using a Gerstel MPS system with 15 μl of 2% (*w*/*v*) methoxyamine hydrochloride (Sigma-Aldrich) in pyridine (incubated for 90 min at 55°C) and 15 μl N-tertbutyldimethylsilyl-N-methyltrifluoroacetamide (MTBSTFA) with 1% tert-butyldimethylchlorosilane (tBDMS) (RESTEK, Cat.# 35601), incubated for 60 min at 55°C. Derivatives were analyzed by gas chromatography (GC) mass spectrometry (MS) using a ZB-35MS column (Phenomenex, Cat.# 7HG-G003-11-GGA-C) installed in an Agilent 7890B gas chromatograph interfaced with an Agilent 5977B MSD mass spectrometer operating under electron impact ionization at 70 eV. The MS source was held at 230 °C, the quadrupole at 150 °C, and helium was used as a carrier gas. The GC oven temperature was held at 100 °C for 2 min, increased to 325 °C at a rate of 10 °C/min, and held at 325 °C for 4 min or increased to 180 °C at a rate of 10 °C/min, followed by a ramp at 5 °C/min to 300°C, with a post-run hold at 325°C for 3 minutes.

The metabolite identification of the GC-MS data was performed by using open chrome open source software (Lablicate) to determine each metabolite’s retention times. Mass isotopomer distributions and total metabolite abundances were computed by integrating mass fragments with corrections for natural isotope abundances using in-house algorithms as described previously^62,63^. Metabolite abundances were normalized to protein content or mg tissue as indicated in the figure legend. Metabolite concentrations were calculated based on external standard curves.

### GC-MS/MS analysis and sample preparation

Sample preparation and derivatization followed the same protocol used for GC-MS analysis. Derivatized samples were analyzed on a high-resolution/accurate-mass (HRAM) Thermo Scientific™ Q Exactive™ GC Orbitrap™ system coupled to gas chromatograph (TRACE 1310) equipped with a ZB-35MS column (Phenomenex, 7HG-G003-11-GGA-C, 30m x 0.25mm x 0.25 µm film). The GC oven temperature was held at 100 °C for 2 min, increased to 180 °C at 10 °C/min and further increased to 300°C at 5°C/min. Helium was used as a carrier gas at a flow rate of 1 ml/min. A programmed temperature vaporization (PTV) injector was employed, initiating at a temperature of 70°C. The injector temperature was increased to 270°C at a rate of 14°C/s and held for 5 min in splitless mode (splitless time 0.8 min). The Orbitrap operated in Electron Ionization (EI) mode (70 eV), scanning a mass range of 50–500 m/z at a resolution of 60,000 (Scan and SIM) or 30,000 (MS/MS) to capture both quantitative and qualitative data. The EI source was held at 200°C and the transferline at 280°C. MS/MS was conducted in full-scan or targeted-SIM mode operated in positive mode, using a normalized collision energy of 30 eV. In targeted-SIM mode, the system was configured with lock masses of m/z = 207.03235, m/z = 281.05114, and m/z = 355.06993 to ensure accurate mass calibration. Additionally, an inclusion list was specified for m/z = 433.22500, 434.22830, 435.22000, 437.23840, and 438.24180, while m/z = 73.05, 75.02, 78.04, and 147.06 were placed on an exclusion list. Multiple reaction monitoring (MRM) was performed by using mass transitions between specific parent ions into corresponding fragment ions for each analyte. Characteristic ions for C5 dicarboxylate compounds are depicted in Supplementary Figures. Data were processed using Thermo Scientific Xcalibur (version 4.3.73.11) and Freestyle software (version 1.6.75.20).

### Liquid chromatography coupled to tandem mass spectrometry (LC-MS/MS) analysis and sample preparation

For the analytical separation of C_5_ dicarboxylate compounds an ultra-high-performance liquid chromatography (UHPLC, Ultimate 3000RS (Dionex/Thermo) coupled to a time-of-flight (TOF) mass-spectrometer (maxis HD, Bruker, Bremen, Germany) was used. The UHPLC was equipped with a Chiral NEA (R) column (100 x 4,6 mm, 5 µm, 30 nm) heated to 40 °C. UHPLC was performed using a gradient of H_2_O with 0.5 % formic acid (solvent A) and 95/5 MeOH/H_2_O with 20 mM ammonium acetate (solvent B) at a flow rate of 900 µl/min. The linear gradient had 0.3 % solvent B at 0 min, 0.3 % solvent B at 3 min, 100 % solvent B at 8 min, and 100 % solvent B at 10 min before returning back to starting condition. The overall run time was 14 minutes. Metabolites were analyzed in MS using an ESI – Apollo II electrospray source with a scan range of 50 – 1500 m/z in negative ion mode, a capillary voltage of 4500 V, nebulizer pressure of 4.0 bar and 9.0 l/min dry gas flow with the dry heater set to 200 °C. Within each run, a sodium formate cluster was infused into the system within the first 0.3 min for internal calibration. For lock mass calibration, an ion source fragment of hexakis(2,2-difluoroethoxy)phosphazene was used additionally (C_10_H_15_F_10_N_3_O_6_P_3_, m/z = 556.0020 for negative ion mode). Calibrations were done with Data Analysis software (Bruker, Bremen, Germany). To perform MS/MS analyses, the collision energy was varied between 17 and 50 eV and stepped from 80 to 200 %, depending on the mass and charge of the parent ion. The collision RF was stepped between 150 and 300 Vpp, and the TOF analyzer’s transfer time was stepped from 20 to 50 µs.

### Synthesis of sodium 2-hydroxymethylsuccinate (2HMS)

Aqueous NaOH solution (prepared by dissolving solid NaOH (Carl Roth, Cat.# 6771.1) in H_2_O) (0.7 ml, 2 M, 1.4 mmol) was added to the solution of 5-oxotetrahydrofuran-3-carboxylate compound (36 mg, 0.28 mmol, BLDpharm, Cat.# BD00824212) in H_2_O (5 ml). The reaction mixture was stirred for 18 h at 40 °C. The reaction mixture was lyophilized yielding a mixture of the desired compound and NaOH. To remove the excess of NaOH the product was purified via a HILIC column (XBridge BEH Amide 5 µm, 10 x 250mm, Waters, Ireland). The solvents were A: 40 mM ammonium acetate in water and B: 100 % acetonitrile. The used gradient was 0 min: 98 % B, 0.5 min: 98 % B, 4.0 min: 78 % B, 28.0 min: 65 % B, 32.0 min: 50 % B, 35.0 min: 50 % B. The product eluted between 25.0 and 29.0 min yielding a mixture of the desired compound and NH_4_OAc (1:0.93) that was used for absolute quantification and biological experiments.

### Nuclear Magnetic Resonance (NMR) methodology

Proton (^1^H), carbon (^13^C) and 2D-nuclear magnetic resonance (NMR) spectra were measured with a Bruker Avance III (500MHz) spectrometer with residual protonated solvent (D_2_O δ 4.79) as standard. The NMR data of the synthesized examples are in agreement with their corresponding structural assignments.

NMR spectra of sodium 2-hydroxymethylsuccinate after lyophilization.

^1^H NMR: (500 MHz, D_2_O + NaOD): δ = 3.66 – 3.54 (m, 2 H), 2.73 – 2.66 (m, 1 H), 2.40 (dd, *J* = 15.2, 6.5 Hz, 1 H), 2.20 (dd, *J* = 15.2, 8.5 Hz, 1 H) ppm.

^13^C NMR: (126 MHz, D_2_O + NaOD): δ = 182.24, 181.22, 63.31, 48.12, 37.56 ppm.

The structure of the product was additionally confirmed by 2D-NMR (^1^H-^1^H COSY, HMBC and HSQC).

### Quantitative real-time polymerase chain reaction (qRT-PCR)

The RNA isolation was performed according to the manufacturer’s instructions for the NucleoSpin^®^ RNA kit (Cat.# 740955.250, Machery-Nagel). The RNA concentration was quantified by using a microplate reader (Tecan Infinite^®^ 200 Spark^®^). 10 µl RNA (100 ng/µl) per sample was reverse transcribed into cDNA using High-Capacity cDNA Reverse Transcriptase kit (Cat.# 4368813, Thermo Fisher Scientific). Individual 10 μl SYBR Green real-time PCR reactions consisted of 2 μl of diluted cDNA, 5 μl iTaq Universal SYBR Green Supermix (Cat.# 1725124, BIO-RAD), and 1.5 μl of each 10 μM forward and reverse primers. For standardization of quantification, *RPL27* was amplified simultaneously. PCR was carried out in 96 well plates (MicroAmp™ Optical 96-Well Reaction Plate, Cat.# N8010560, Applied Biosystems by Life technologies^®^) and centrifuged at 300 x *g* for 30 sec before measured with QuantStudio Design & Analysis Software v1.5.2 (Thermo Fisher Scientific) using a three-stage program provided by the manufacturer: 95 °C for 20 sec, 45 cycles of 95 °C for 1 sec and 60 °C for 20 sec.

Primer uses are as follows. *Auh* (mouse) forward: GCATTCCAGTGAAGTTGGTCCC; *Auh* (mouse) reverse: ATGTCACACGCTAGAGCCAGCT; *Irg1* (mouse) forward: GCAACATGATGCTCAAGTCTG; *Irg1* (mouse) reverse: TGCTCCTCCGAATGATACCA; *Rpl27* (mouse) forward: GCGATCCAAGATCAAGTCCTTTG; *Rpl27* (mouse) reverse: TCAAAGCTGGGTCCCTGAACAC; *Slc13a3* (mouse) forward: CAGGGAAATGGACTGCGAACAG; *Slc13a3* (mouse) reverse: TCCTCCTTGGAGTCATCAGGCA; *Suclg1* (mouse) forward: GTCTTACACAGCCTCTCGGAAAC; *Suclg1* (mouse) reverse: ACTCCAAAGCCTGCTGACTGTG; *Sugct* (mouse) forward: CAGGACCCTTTGCCACGATGAA; *Sugct* (mouse) reverse: CCAAGATCGTGTGTCATCACCAG.

### Gene silencing

For siRNA-mediated silencing, ON-TARGETplus SMARTpool siRNAs targeting murine *Suclg1* (gene ID: 56451, L-049787-01), *Auh* (gene ID: 11992, L-049078-01), and *Sugct* (gene ID: 192136, L-042443-01) were obtained from Horizon Discovery. A non-targeting siRNA pool (siCtr, Cat.# D-001810-10) was used as the control. Each siRNA was reconstituted in RNase-free water to a stock concentration of 20 µM by dissolving 5 nmol of siRNA in 250 µl. RAW264.7 murine macrophage-like cells were electroporated using the Amaxa SF Cell Line 4D-Nucleofector X kit L (Cat.# V4XC-2024, Lonza) with the 4D-Nucleofector system (Cat.# AAF-1003, Lonza). Briefly, 2 million cells were resuspended in 100 µl of SF Nucleofector Solution mixed with 60 pmol of the appropriate siRNA. Electroporation was performed using the program code optimized for RAW246.7 cells. After electroporation, cells were incubated for 5 minutes at room temperature within the Nucleocuvette. Subsequently, 500 µl pre-warmed complete DMEM was added to each cuvette, and the contents were gently mixed by pipetting up and down. The cell suspension was then plated into 12-well plates at a density of 0.5 million cells per well and incubated for 24 h. Following incubations, cells were treated with 10 ng/ml LPS, 10 mM itaconate or 10 mM mesaconate for 24 h before metabolite extraction and RNA isolation.

3T3-L1 pre-adipocytes were seeded at a density of 120,000 cells per well in 12-well plates and cultured overnight in DMEM supplemented with 10 % FBS and penicillin-streptomycin. For each well, siRNA-Lipofectamine complexes were prepared by diluting 50 pmol of siRNA in 400 µl Opti-MEM I Reduced-Serum Medium (Cat.# 31985-062, Invitrogen). Separately, 2 µl of Lipofectamine RNiMAX transfection Reagent (Cat.# 13778100, Invitrogen) was added to the diluted siRNA solution, mixed gently, and incubated at room temperature for 20 min to allow complex formation. The culture medium was removed from the cells, and 400 µl of the siRNA-lipid complexes was added directly to each well. Cells were incubated with the transfection complexes for 6 h under standard culture conditions. Subsequently, 1 ml of complete DMEM medium was added to each well without removing the transfection mixture, and cells were incubated for an additional 18 h. Following the 24 h transfection period, cells were treated with 10 mM itaconate for 24 h prior to metabolite extraction and RNA isolation.

### RNA-seq experiment and data analysis

RAW264.7 cells were treated with 500 µM 2HMS or 465 µM ammonium acetate as control condition (1:93 molar ratio relative to 2HMS) for 4 h before stimulation with 10 ng/ml LPS for 3 h. Total RNA was isolated according to the manufacturer’s instructions using the NucleoSpin® RNA Isolation Kit (Macherey-Nagel, Cat.#740955.250). The quality and integrity of total RNA were assessed using the Agilent Fragment Analyzer 5200 (Agilent Technologies, Waldbronn, Germany). RNA sequencing libraries were prepared from 250 ng of total RNA with the NEBNext® Ultra™ II Directional RNA Library Prep Kit for Illumina®, including the NEBNext Poly(A) mRNA Magnetic Isolation Module (New England BioLabs), following the manufacturer’s instructions. Libraries were sequenced on an Illumina NovaSeq X Plus platform using an Illumina 1.5 Flowcell (300 cycles, paired-end) with an average yield of 3 × 10⁷ reads per RNA sample. Data analysis was performed using the nf-core/rnaseq pipeline (version 3.19.0)^64^, followed by the nf-core/differential abundance pipeline (version 1.5.0)^65^. Data processing of differentially expressed genes was conducted in *R* (version 4.3.0)^66^.

Adju^64^sted *p*-values (padj) were transformed into –log_10_ values. Genes were classified into three categories: *up-regulated* (padj < 0.05, log_2_FC > 0), *down-regulated* (padj < 0.05, log_2_FC < 0), and *non-significant*. The ten most significant genes per category were highlighted based on the padj. Data visualization was performed with ggplot2 (version 3.5.2)^67^ and ggrepel (version 0.9.6)^68^. Volcano plots display log_2_FC and –log_10_(padj). Thresholds were indicated by dashed lines at log_2_FC ±0.5 and padj = 0.05.

### *In vivo* ^13^C itaconate mice study

In our previous study, we performed ^13^C itaconate *in vivo* experiments with male mice and generated metabolic data using GC-MS^28^. For the current study, we re-analyzed the data for 2HMS integration. ^13^C itaconate was provided by NIH Common Fund Metabolite Standards Synthesis Core (NHLBI Contract No. HHSN268201300022C). Briefly, C57BL/6J mice were administered 400 mg/kg body weight itaconate via retroorbital injection. Plasma samples were collected at indicated time points. Tissues were collected from *n* = 5 animals for 15 min ^13^C itaconate and *n* = 4 for 45 min ^13^C itaconate treatment. Data were compared to control condition (0 min itaconate), where NaCl was given for 45 min with *n* = 4 animals.

### CpG oligodeoxynucleotides (ODNs) *in vivo* model

The animal study was approved by the Intramural Committee for Animal Experimentation at the Medical University of Vienna and the Austrian Ethics Committee for Animal Experiments (GZ BMBWF-2023-0.380.996). Male C57BL/6J mice at the age of 14 weeks were purchased from Janvier Labs and were housed in the Mouse Vivarium at the Institute for Medical Genetics at the Medical University of Vienna. Mice were acclimated to the facility for at least 7 days prior to any experimental manipulation. The housing facility was maintained on a 12-hour light/dark cycle with ad libitum access to chow (Altromin, Cat.#1324) and autoclaved water. The CpG oligonucleotide (ODN1826 sodium salt, MCE, Cat.# HY-1416245C) was dissolved in dPBS to a concentration of 1 mg/ml and filtered using a 0.2 µm PTFE syringe filter (Fisher Scientific, Cat.# 15342378). Within each cage, mice were randomly assigned to the control (dPBS, *n* = 5) or CpG treatment groups (*n* = 5). Mice received a 200 µl intraperitoneal injection (corresponding to 200 µg CpG) every second day for a total of 5 injections (Fig. 5a). The injection site was alternated between the left and right side at each administration. All blood and tissue collection was performed two days after the final injection. Mice were fasted for eight hours prior to sample collection. Blood was collected by tail bleed from awake, non-restrained mice into lithium heparin-coated capillary collection tubes (Sarstedt, Cat.# 16.443.100), placed on ice, and immediately spun in a refrigerated microcentrifuge at 15,000 x *g* for 5 min at 4 °C. Plasma was then transferred to fresh polypropylene tubes on dry ice. Following blood sampling, mice were euthanized by cervical dislocation and their tissues rapidly dissected. Solid tissue was wrapped in foil and clamped using Wollenberger tongs pre-cooled in liquid nitrogen to immediately quench metabolism throughout the sample. Bone marrow was collected via centrifugation as described^69^ and snap-frozen in liquid nitrogen. Abundances of 2HMS in tissue and plasma samples were calculated with the fragment m/z 433 and subtraction of an overlaying background peak (m/z 432, C_19_H_42_O_4_N_1_Si_3_).

### Statistics

Data visualization and statistical analysis were performed using GraphPad Prism (v10.3.1, GraphPad Software), Adobe Illustrator CS6 (v24.1.2, Adobe Inc.), ChemDraw (v23.1.1), and Biorender.com. The type and number of replicates and the statistical test used are described in each figure legend. Data are presented as means ± standard error of mean (s.e.m.) or box (25th–75th percentile with median line) and whiskers (minimum to maximum values). Experiments were independently repeated as indicated in the figure legend. *P* values were calculated using an unpaired *t*-test, one-way ANOVA or two-way ANOVA with Fisher’s least significant difference (LSD) post hoc test. For all tests, *p* < 0.05 was considered significant with \**p* < 0.05; \*\**p* < 0.01; \*\*\**p* < 0.001, and ^#^*p* < 0.0001 as indicated in each figure legend.

## Availability of data

All the data associated with the study are in the paper or Supplementary Data. Source data will be deposited in the repository platform of Technische Universität Braunschweig with a dedicated DOI upon acceptance of this manuscript.

## Acknowledgement

We thank all members of the Cordes Lab for helpful discussions. This study was supported in part by internal funds from Technische Universität Braunschweig (to T.C.) and the Helmholtz Centre for Infection Research (to T.C.), by an Exploration Grant of the Boehringer Ingelheim Foundation (BIS) (to T.C.), by the Austrian Science Fund (FWF) grants 10.55776/P34023, 10.55776/P34266, FWF Sonderforschungsbereich F83 (10.55776/F83) (to T.W.), and by the Ann Theodore Foundation Breakthrough Sarcoidosis Initiative (to T.W.). We thank Sabine Kaltenhäuser for her technical expertise and support with mass spectrometry measurements. We thank the NIH Common Fund Metabolite Standards Synthesis Core (NHLBI Contract No. HHSN268201300022C) for providing isotopic labeled [U-^13^C_5_]itaconate. We acknowledge support from the Open Access Publication Funds.

## Contributions

Conceptualization, F.C., T.C.; Methodology, F.C., L.H., V.S.K., U.B., C.S.R.J., R.F., T.C.; Formal analysis, F.C., L.H., T.C.; Investigation, F.C., L.H., B.D., V.S.K., U.Be., C.S.R.J., H.F.W., U.Br., R.F., M.B., M.N-S, T.C.; Resources: E.M., M.J., H.G., A.F., T.W., M.B., M.N-S, T.C.; Writing – original draft, T.C. with the help of F.C.; Writing – reviewing and editing, all authors; Visualization, F.C., L.H., U.B., V.S.K., T.C.; Project administration, T.C.; Supervision, T.C.; Funding acquisition, T.W., T.C. All authors have read and agreed to the published version of the manuscript.

## Supplemental information

Document S1, Figures S1 – S6

## Abbreviations

ACOD1: *cis*-Aconitate Decarboxylase (protein)
AT: Adipose Tissue
ATCC: American Type Culture Collection
Auh: AU RNA Binding Methylglutaconyl-CoA hydratase (gene)
AUH: Methylglutaconyl-CoA hydratase (protein), also known as 3-MGH
B_12_: Vitamin B_12_ (Cobalamin)
BAT: Brown Adipose Tissue
BCAA: Branched-chain amino acid
BMDM: Bone Marrow-Derived Macrophages
12C: Carbon-12
13C: Carbon-13
CA: Citraconate
CDC: CoA-dependent conversion
CIC: CoA-independent conversion
CiMa: Citramalate
CoA: Coenzyme A
CpG ODN: Cytosine-phosphorothioate-guanine oligodeoxynucleotides
Ctr: Control
DMEM: Dulbecco’s Modified Eagle’s Medium
FBS: Fetal Bovine Serum
FELASA: Federation of European Laboratory Animal Science Associations
GC-MS: Gas Chromatography–Mass Spectrometry
GC-MS/MS: Gas Chromatography-Tandem Mass Spectrometry
gWAT: Gonadal White Adipose Tissue
2HG: 2-Hydroxyglutarate
3HG: 3-Hydroxyglutarate
2HMS: 2-Hydroxymethylsuccinate
hMDM: Human Monocyte-Derived Macrophages
IFNγ: Interferon-gamma
*Irg1*: Immunoresponsive gene 1 (gene)
IRG1: Immune-Responsive Gene 1 Protein
ita: Itaconate
KO: Knockout
LC-MS/MS: Liquid Chromatography-Tandem Mass Spectrometry
LPS: Lipopolysaccharide
3MeMa: 3-Methylmalate
mesa: Mesaconate
MGCA1: 3-methylglutaconic aciduria type 1
MMA: Methylmalonate
MTBSTFA: N-tert-Butyldimethylsilyl-N-methyltrifluoracetamid
MUT: Methylmalonyl-CoA mutase (protein)
m/z: Mass-to-charge ratio
NaDC3, SLC13A3: Sodium-dependent dicarboxylate transporter 3 (protein)
NMR: Nuclear Magnetic Resonance
PBMC: Peripheral Blood Mononuclear Cells
rPFO: Recombinant perfringolysin
RPMI: Roswell Park Memorial Institute medium
RT: Retention time
SA: Standard addition
SCS: Succinyl-CoA synthetase (protein)
scWAT: Subcutaneous White Adipose Tissue
s.e.m.: Standard error of mean
siRNA: Small interfering RNA
*Slc13a3*: Solute carrier family 13 member 3 (gene)
*Suclg1*: Succinate-CoA ligase GDP/ADP-forming subunit alpha (gene)
*Sugct*: Succinyl-CoA:glutarate CoA-transferase (gene)
SUGCT: Succinyl-CoA:glutarate CoA-transferase (protein)
TCA: Tricarboxylic acid cycle
tBDMS: tert-Butyldimethylsilylchlorid
TIC: Total ion chromatogram
TLR: Toll-like receptor
U: Uniformly
[U-^13^C]substrate: Uniformly labeled ^13^C carbon tracer
WAT: White Adipose Tissue
WT: Wild-Type

---

**Supplementary Figure S1:**
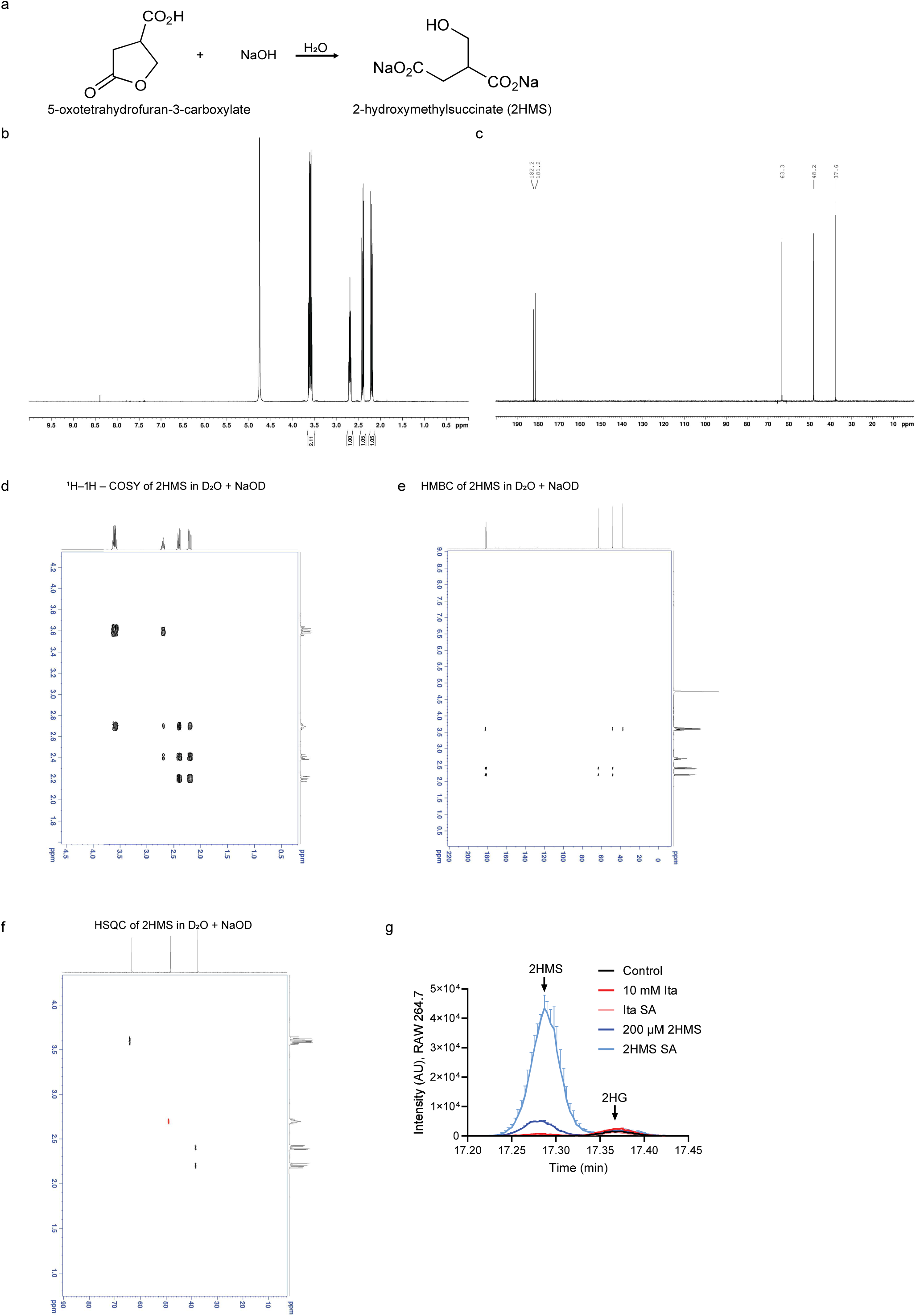
Synthesis and analytical characterization of a 2HMS standard. **a,** Alkaline hydrolysis of 5-oxotetrahydrofuran-3-carboxylate for sodium 2-hydroxymethylsuccinate (2HMS) synthesis. **b,** ^1^H-NMR spectrum and **c,** ^13^C-NMR spectrum of the synthesized compound sodium 2-hydroxymethylsuccinate in D^2^O + NaOD. **d,** ^1^H–^1^H – COSY of sodium 2-hydroxymethylsuccinate in D_2_O + NaOD. **e,** HMBC of sodium 2-hydroxymethylsuccinate in D_2_O + NaOD1. **f,** HSQC of sodium 2-hydroxymethylsuccinate in D_2_O + NaODH. **g,** Single ion (m/z 433) chromatogram depicting 2-hydroxymethylsuccinate (2HMS), and 2-hydroxyglutarate (2HG) in RAW264.7 cells upon standard addition (SA) or cultured with 10 mM itaconate and 200 µM 2HMS for 24 h. Data in g. are presented as mean ± s.e.m. from three cellular replicates.

**Supplementary Figure S2:**
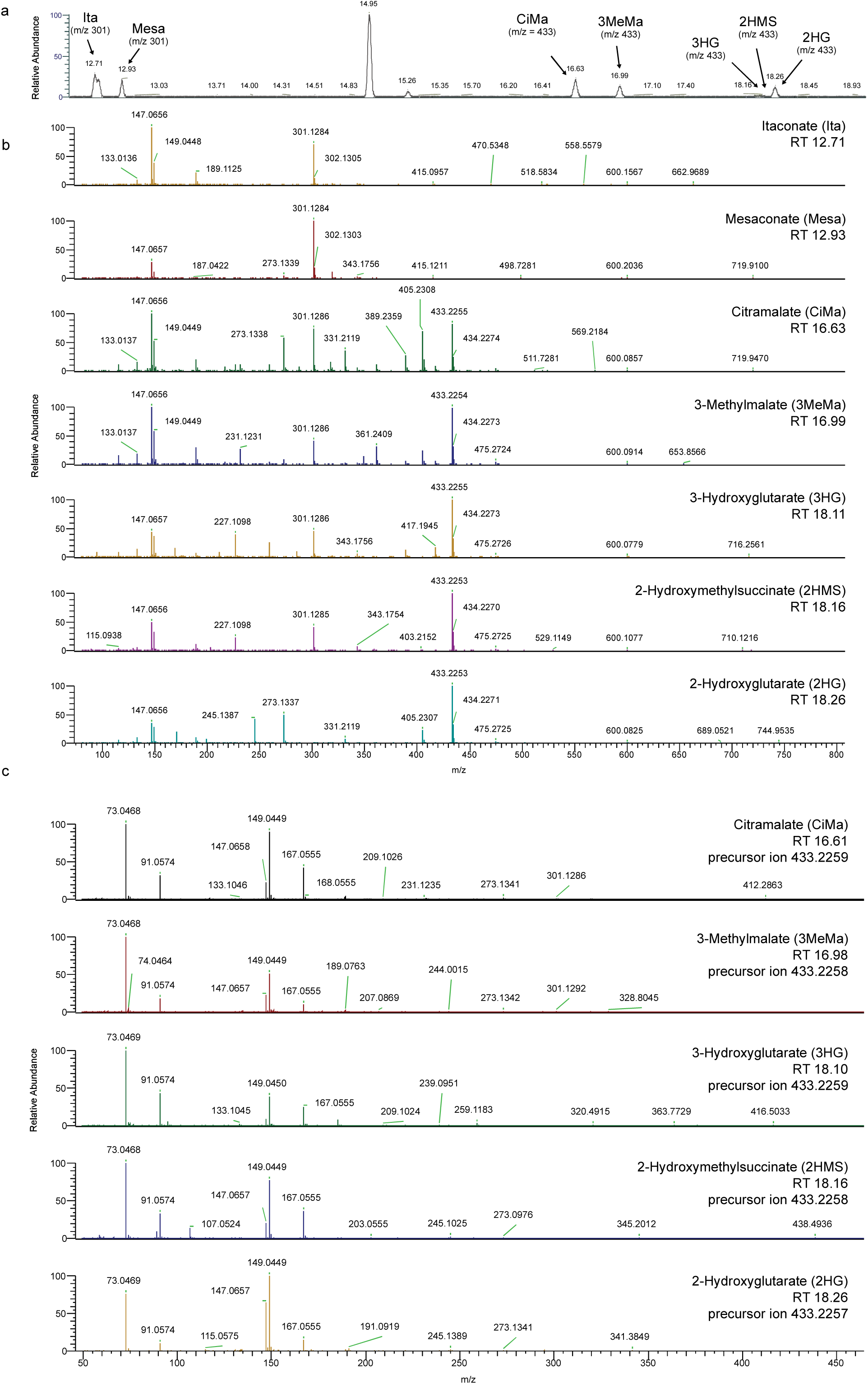
Chromatograms and high-resolution mass spectra of C_5_ dicarboxylate compounds using GC-MS/MS. **a,** Total Ion Chromatograms (TIC) of synthetic standards of C_5_ dicarboxylate compounds separated by retention time (RT). **b,** High-resolution mass spectra for C_5_ dicarboxylate compounds and their characteristic MS1 mass fragments. **c,** High-resolution mass spectra for C_5_ dicarboxylate compounds and their characteristic MS/MS mass fragments (product ions) obtained from precursor ion m/z 433.2259.

**Supplementary Figure S3:**
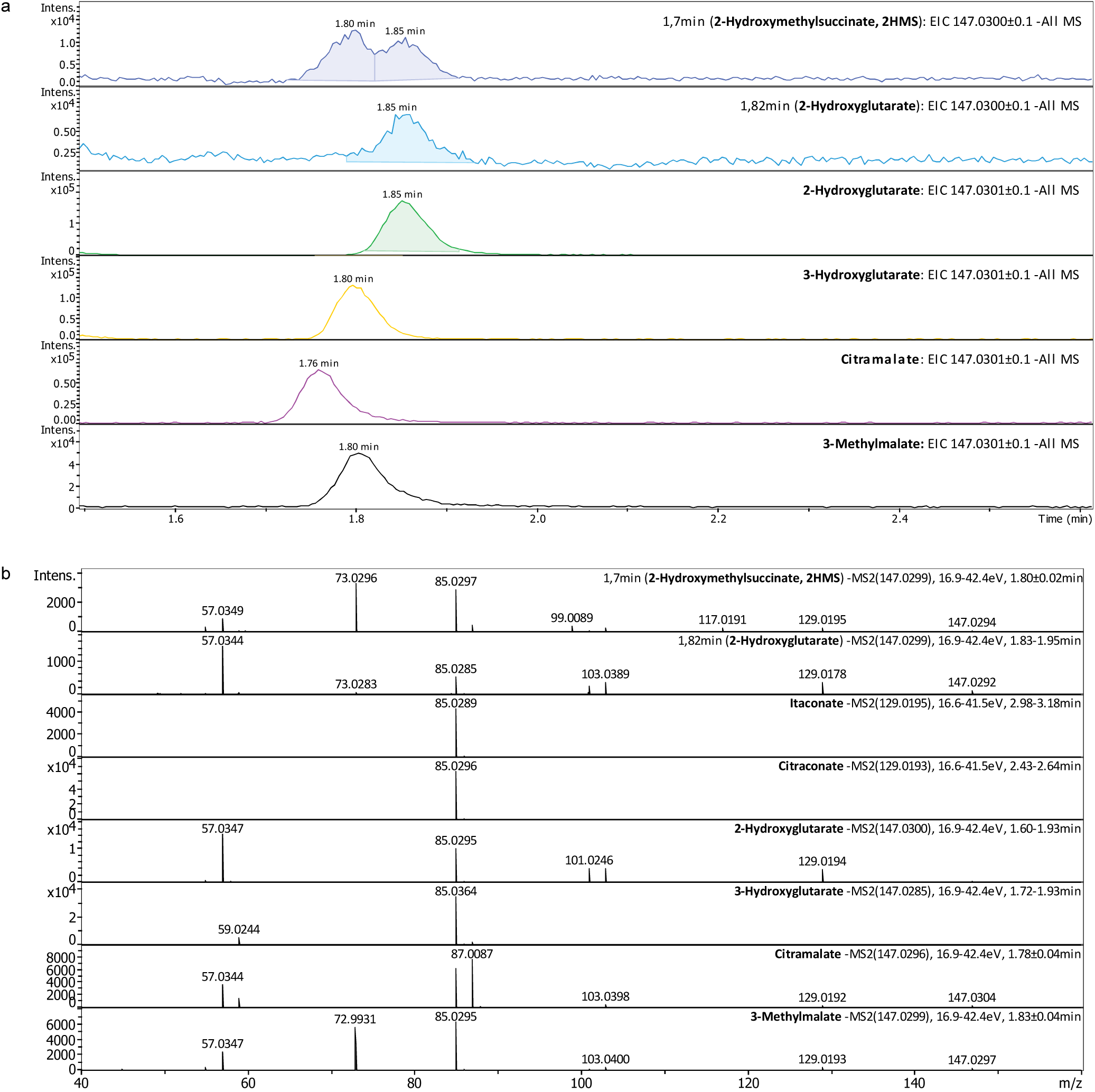
Chromatograms and high-resolution mass spectra of C_5_ dicarboxylate compounds using LC-MS/MS. **a,** Total Ion Chromatogram of C_5_ dicarboxylate compounds. Cellular metabolite fractions eluted at 1.7 min (containing 2HMS) and 1.82 min (containing 2HG) were obtained from RAW264.7 macrophages cultured for 24 h with 10 mM itaconate. Other compounds depict synthetic standards. **b,** High-resolution mass spectra for cell substrates and C_5_ dicarboxylate compound standards and their characteristic MS/MS mass fragments (product ions) obtained from precursor ion m/z 147.0300 (2HMS, 2HG, 3HG, citramalate, and 3-methylmalate) or m/z 129.0195 (itaconate, citraconate).

**Supplementary Figure S4:**
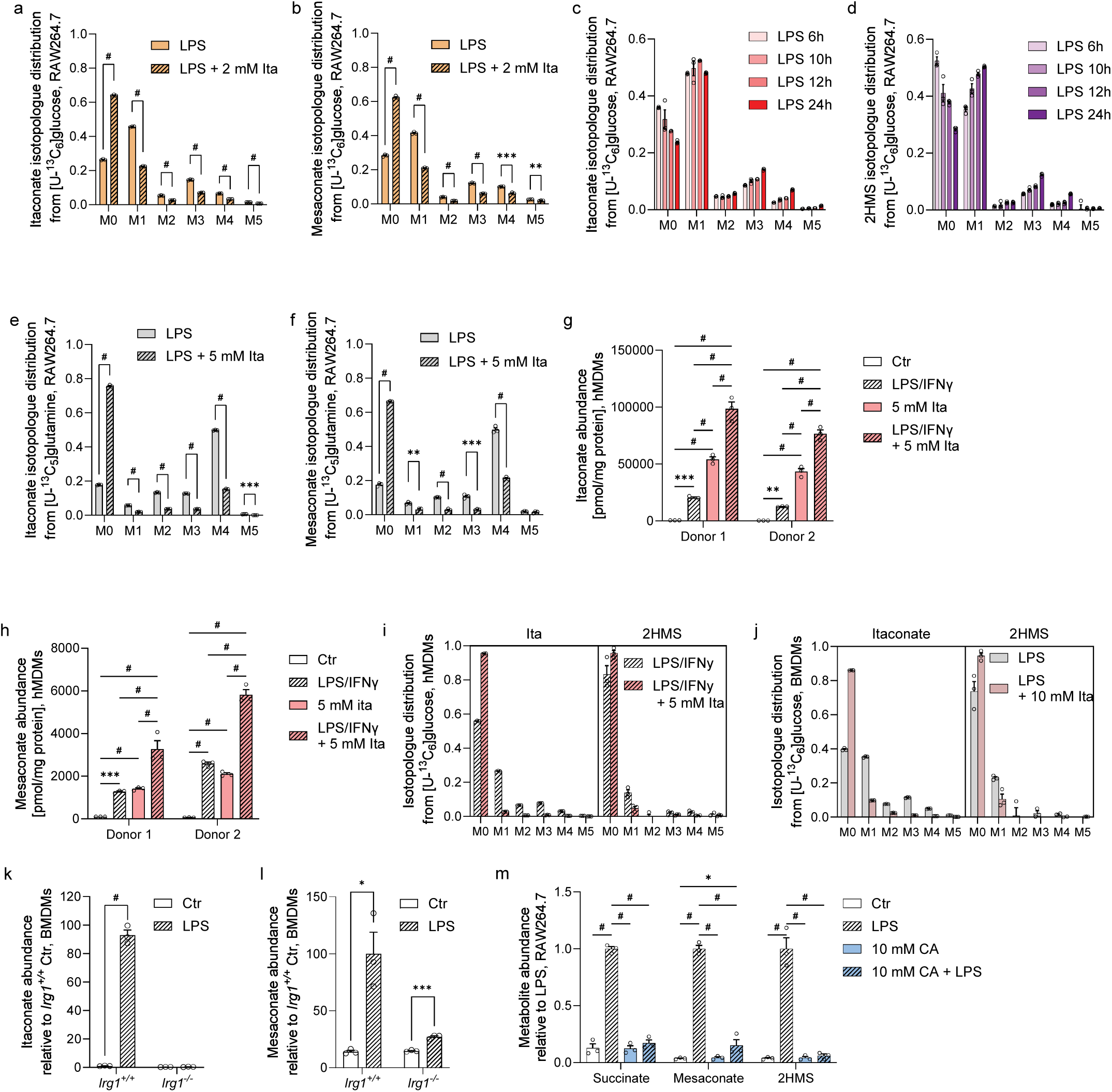
Synthesis of 2-hydroxymethylsuccinate (2HMS) in macrophages. **a,** Itaconate and **b,** Mesaconate isotopologue distributions in RAW264.7 cells cultured for 24 h with 2 mM itaconate and 10 ng/ml LPS from [U-^13^C_6_]glucose. **c**, Itaconate and **d**, 2HMS isotopologue distributions in RAW264.7 cells cultured with [U-^13^C_6_]glucose and 10 ng/ml LPS for 6, 10, 12, and 24 h. **e**, Itaconate and **f**, Mesaconate isotopologue distributions in RAW264.7 cells cultured with 5mM itaconate and 10 ng/ml LPS from [U-^13^C_5_]glutamine. **g**, Itaconate and **h**, Mesaconate abundances in monocyte-derived macrophages (hMDMs) cultured for 24 h with 10 ng/ml LPS, 400 U/ml interferon y (IFNy), and 5 mM itaconate. **i**, Itaconate and 2HMS isotopologue distributions in hMDMs cultured with [U-^13^C_6_]glucose, 100 ng/ml LPS, 400 U/ml IFNy, and with 0 or 5 mM itaconate for 24 h. **j**, Itaconate and 2HMS isotopologue distribution in bone marrow-derived macrophages (BMDMs) cultured with [U-^13^C_6_]glucose and 10 ng/ml LPS with or without 5 mM itaconate for 24 h. **k**, Itaconate and **l,** mesaconate abundance in BMDMs obtained from immune responsive gene 1 deficient (*Irg1*^-/-^) and wildtype (*Irg1*^+/+^) mice and cultured for 24 h with 10 ng/ml LPS. **m**, Metabolite abundance in RAW264.7 cells cultured with 10 mM citraconate (CA) and 10 ng/ml LPS for 12 h. Data are presented as means ± s.e.m. with three cellular replicates. Each experiment was repeated independently three times (a-b, e-f), two times (c, d, m), or with *n* = 3 mice (j, k, l) and three donors (g, h, i). *P* values were calculated by multiple unpaired *t*-test (a, b, e, f), two-way ANOVA (g, h, k-m) with * *p* < 0.05; ** *p* < 0.01, *** *p* < 0.001, ^#^ *p* < 0.0001.

**Supplementary Figure S5:**
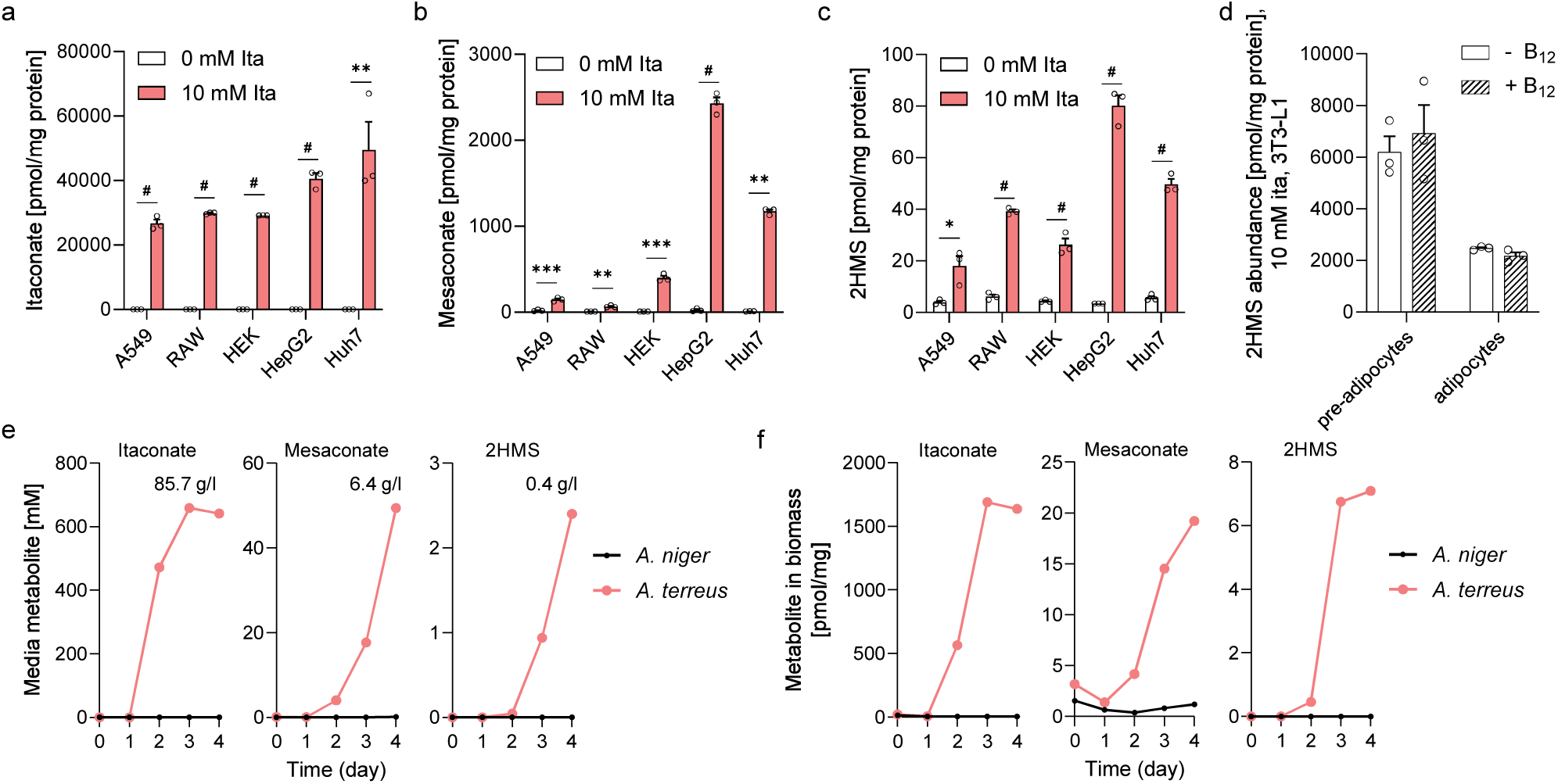
2-Hydroxymethylsuccinate (2HMS) synthesis is cell-type specific. **a,** Itaconate abundance in different cell types cultured with 0 mM or 10 mM itaconate for 12 h. **b,** Mesaconate abundance in different cell types cultured with 0 mM or 10 mM itaconate for 12 h. **c,** 2HMS abundance in different cell types cultured with 0 mM or 10 mM itaconate for 12 h. **d,** 2HMS abundance in pre- and adipocytes cultured with 1 µM vitamin B_12_ for 24 h. **e**, Media or **f,** biomass metabolite abundances in *Aspergillus terreus* and *Aspergillus niger* in a culture of 4 days and calculated yields (g/l) at day 4. Data are presented as means ± s.e.m. with three cellular replicates (a-d). Each experiment was repeated independently three times (d) or two times (a-c) with similar results. *P* values were calculated by multiple unpaired *t*-test with * *p* < 0.05; ** *p* < 0.01, *** *p* < 0.001, ^#^ *p* < 0.0001.

**Supplementary Figure S6:**
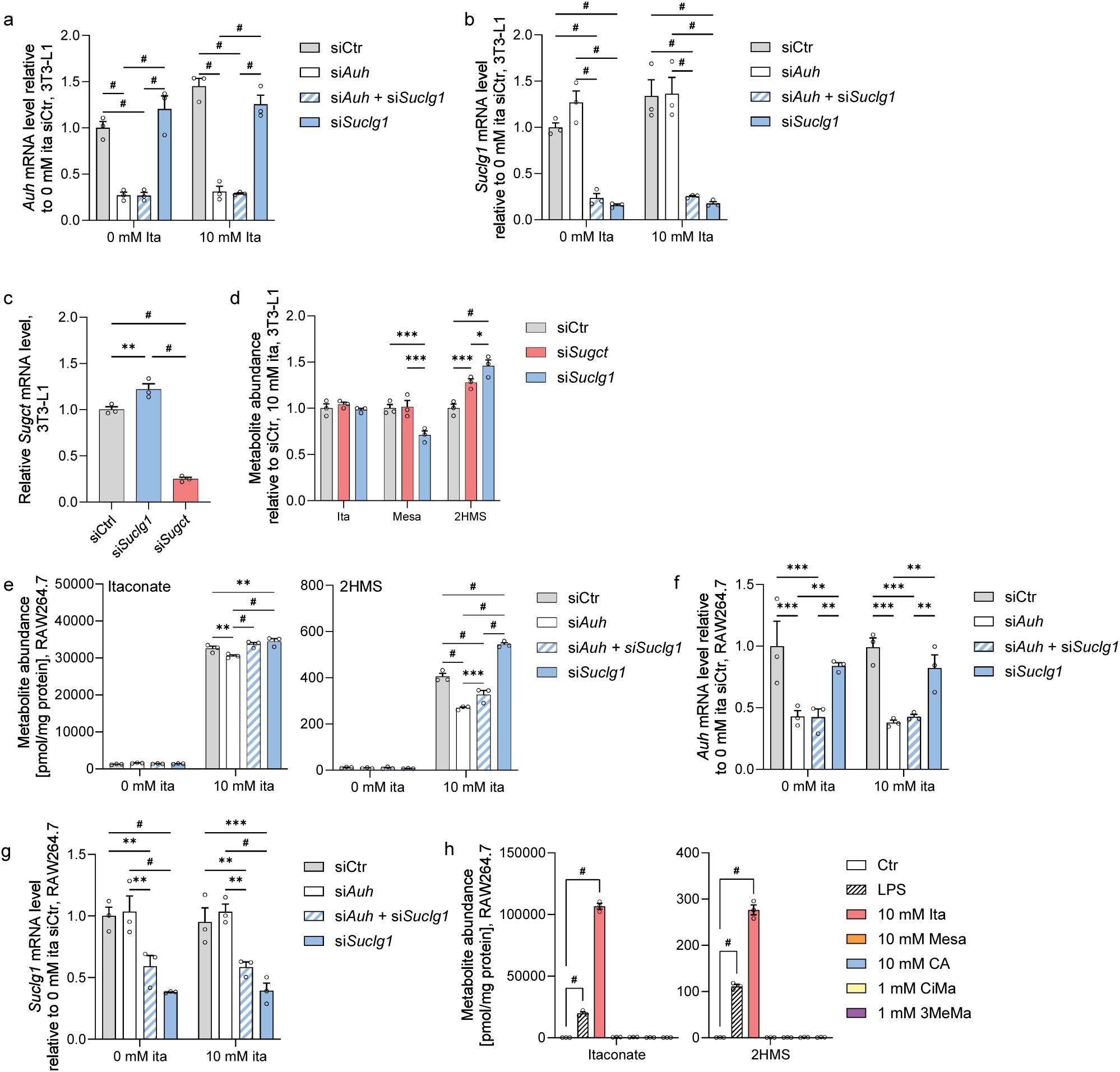
Itaconate degradation pathway for 2-hydroxymethylsuccinate (2HMS) and mesaconate synthesis. **a,** Methylglutaconyl-CoA hydratase (*Auh*) expression level in 3T3-L1 cells with small interfering RNA (siRNA)-mediated gene knockdown targeting *Auh*, succinate-CoA ligase (*Suclg1*), and both genes combined (si*Auh* + si*Suclg1*) compared to control condition (siCtr) cultured with 0 mM or 10 mM itaconate for 24 h. **b,** *Suclg1 expression* level in 3T3-L1 cells in response to siRNA-mediated gene knockdown of *Auh* and *Suclg1* cultured with 0 mM or 10 mM itaconate for 24 h. **c,** Succinyl-CoA:glutarate CoA-transferase (*Sugct1*) expression level in 3T3-L1 cells. **d,** Metabolite abundance in 3T3-L1 cells in response to siRNA-mediated knockdown of *Sugct* and *Suclg1*. **e,** Metabolite abundance in RAW264.7 cells in response to siRNA-mediated gene knockdown of *Auh* and *Suclg1,* and both genes combined (*siAuh* + *siSuclg1*), cultured with 0 mM or 10 mM itaconate for 24 h. **f**, *Auh* expression level in RAW264.7 cells with siRNA-mediated gene knockdown targeting *Auh* and *Suclg1* cultured with 0 mM or 10 mM itaconate for 24 h. **g,** *Suclg1 expression* level in RAW264.7 cells in response to siRNA-mediated gene knockdown of *Auh* and *Suclg1* cultured with 0 mM or 10 mM itaconate for 24 h. **h,** Metabolite abundance in RAW264.7 cells cultured for 24 h with itaconate (ita), mesaconate (mesa), citraconate (CA), citramalate (CiMa) or 3-methylmalate (3MeMa) compared to untreated control cells (Ctr). Data are presented as means ± s.e.m. with three cellular replicates. Each experiment was repeated independently three (a, b) or two (c-h) times with similar results. *P* values were calculated by two-way ANOVA (a, b, d-g) or one-way ANOVA (c, h, with h compared to Ctr condition) with * *p* < 0.05; ** *p* < 0.01, *** *p* < 0.001, ^#^ *p* < 0.0001.

